# An explainable machine learning consensus framework for robust estimations of environmental effects on population dynamics

**DOI:** 10.64898/2026.05.10.724190

**Authors:** Anuradha Dhananjanie, Helen Thompson, Julie Vercelloni, David J. Warne

## Abstract

Explainable machine learning (ML) methods are gaining increasing attention in environmental and ecological research for their ability to reveal relationships between environmental drivers and population dynamics. However, there remain questions on the reliability of these tools, especially given recent research shows that these explanations can be highly sensitive to model architecture. In ecology, it is typical to use a single ML model, and a comparative evaluation of sensitivity of explainability for different ML approaches is overlooked. In this paper, we develop a novel framework that quantifies explanation consistency between multiple ML model architectures. This framework provides a discrepancy measure for each model prediction, with high discrepancy indicating substantive explanation disagreement across models and low discrepancy indicating strong consensus in explanations across models. We then demonstrate that low explanation discrepancy aligns well with ground truth mechanism. Furthermore, high explanation discrepancy provide a mechanism to identify areas for model refinement and further investigation by domain experts. We do this by using a simulation study based on synthetic coral cover data that incorporate spatio-temporal variability driven by known disturbance effects. Our method provides a quantitative approach to assess the sensitivity of explainable ML in the absence of ground truth. As a result, this enhances the utility of ML approaches in conservation and ecological management. While we focus primarily on ecological modelling for coral reefs, our methods are generally applicable to other ecological and environmental modelling settings.

## Introduction

With the increasing rate of species extinction caused by human activities such as anthropogenic climate change, overhunting, and species translocations and introductions, conservation of ecosystems has become more crucial than ever (Cowie et al., 2022; Lande, 1998). Models informed by ecological data play a vital role in planning and implementing conservation and management activities. These ecological models are widely used to predict species abundance across space and time, and to identify key environmental factors influencing their dynamics (Hao et al., 2019; Vasconcelos et al., 2024; Ryo et al.,2021; Mellin et al., 2010; El Assari et al., 2023). The importance of ecological models for conservation planning and decision making has led to a substantial body of literature from the ecological modelling community, encompassing a wide range of mathematical and statistical modelling approaches (Beery et al., 2021; Lehmann et al., 2003; Laxton et al., 2023; Akinlotan et al., 2024; Cure et al., 2024; Cattoni et al., 2024).

In recent years, machine learning (ML) approaches have become increasingly popular for ecological modelling, as they are often reported to provide high predictive accuracy, identify spatial patterns, and capture complex trends in data (Makridakis et al., 2018; van den Hoogen et al., 2021; Jelovica et al., 2024; Vulova et al., 2025; Ryo et al., 2021; Forrest et al., 2026). ML techniques have improved the predictive performance of ecological models by enabling the development of more complex models that capture non-linear relationships between features and responses (Athni et al., 2024). However, unlike statistical or mechanistic models, a major challenge in the use of ML is its “black-box” nature (Zhang et al., 2021), which makes these approaches less suitable for identifying the drivers of population changes or extracting ecological insights (Elith and Leathwick, 2009; Yu et al., 2021; Laxton et al., 2023). Therefore, despite their growing popularity in ecological modelling, ML techniques have often been used primarily for predictive purposes rather than for understanding underlying ecological mechanisms (Lucas, 2020; van den Hoogen et al., 2021).

Moreover, the advances in ML algorithms in terms of model accuracy have not necessarily increased our understanding of how these models generate predictions (Ryo et al., 2021). From an ML perspective, understanding the predictive process, results, and relative importance of variables are collectively referred to as model explainability (El Assari et al., 2023). There is a general agreement within the literature that there is a trade-off between model predictive accuracy and explainability (Herm et al., 2023; Burkart and Huber, 2021; Breiman, 2001b; Rudin, 2019; Sanderson et al., 2023). Achieving both high accuracy and explainability simultaneously remains challenging (Ryo et al., 2021). To inform management and decision-makers, modelling must provide more than predictive performance. In this context, models need to be interpretable to enable ecologists to inspect and analyse them, and to confidently identify the most important factors and their effect sizes (Beery et al., 2021). As results from ecological models can directly inform conservation and management decision making, the ability to interpret how a model generates predictions is critical (Ryo et al., 2021). That is, both predictive performance and explainability of an ecological model are equally important. Improving the explainability of ML approaches is therefore crucial for ensuring reliable model interpretation (Hui et al., 2022) and for supporting well-informed conservation and management decisions (Angelov, 2019).

To address this explainability problem in ML models, a range of post-hoc methods for interpreting model predictions exists within the emerging field of explainable ML. These approaches aim to improve understanding of how “black-box” models generate predictions and to provide insight into the patterns learned from data (Molnar, 2019). Existing studies that explore explainable ML in ecological applications (Ryo et al., 2021; El Assari et al., 2023; Mengjuan et al., 2025; Angelov, 2019; Vulova et al., 2025; Cheung et al., 2025; Bai et al., 2025) have typically considered a single ML model, and a comparative evaluation of different ML approaches with respect to their explanatory behaviour does not exist in the ecological literature. However, in the ML and decision support literature, it has been shown that different machine learning models can lead to different explanations (Gwinner et al., 2024; Agarwal et al., 2022). These studies do not highlight an explicit method for identifying extreme discrepancies and do not test the consistency or meaningfulness of explanations. These findings have serious implications as there is limited work that questions how explanations are interpreted in practice within an ecological context. Importantly, different ML models can produce inconsistent explanations for the same observation, even when their predictive performance is similar. This raises the question of how these methods can be used in practice to confidently determine environmental effects on population dynamics under unknown ground truth.

Figure 1 illustrates this issue by comparing explanations from two ML models for an identical input, where substantial differences are observed in both the direction and magnitude of feature contributions (Figure 1(d)–(e)). Such discrepancies raise critical questions regarding which explanations should be trusted and how conflicting explanations should be handled when identifying drivers of ecosystem decline. Currently, no principled framework exists to systematically evaluate the reliability or consistency of explanations, nor to address explanation discrepancies in practical ecological applications. Motivated by this research gap, we propose a framework for evaluating and comparing explanations across multiple ML models. We then investigate how informative explainable ML methods are for identification of associations between environmental drivers and population dynamics. Using a simulation-based study in a marine ecology context, where data are generated from an expert-informed ecological model, we are able to examine explanation behaviour under a controlled data-generating process. This enables us to assess not only whether explanations differ across models, but also whether they are potentially misleading relative to the underlying process used to generate the data.

**Figure 1.**
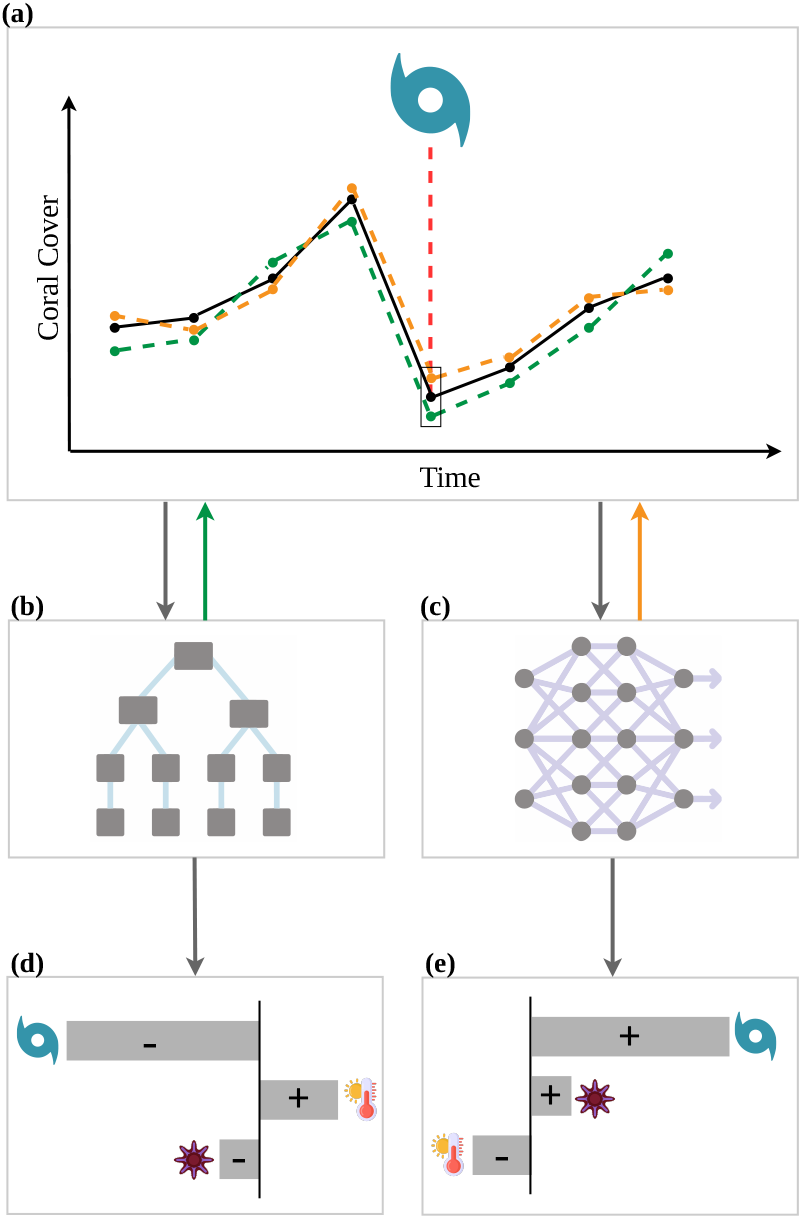
Schematic of different explanations provided by two ML models for the same observation.(a) shows a hypothetical scenario in which a severe storm disturbance event causes a decline in coral cover (black line represents the observed coral cover over the time). A (b) tree-based method and an (c) artificial neural network are used to model the coral cover and are equivalent in predictive performance (green and orange dashed lines represent predictions from the models b and c, respectively.) (d) and (e) illustrate the explanations given as the contribution of features (e.g., storms, temperature and crown-of-thorns starfish) to the model prediction at the time point of coral decline by models (b) and (c), respectively. The sign indicates the impact on the prediction by features, whether it is positive or negative. The length of horizontal bars represents the magnitude of that impact.

Using a state-of-the-art explainable ML method, we conduct a comparative analysis to examine the sensitivity of both local and global explanations across multiple ML approaches. We demonstrate that models with similar predictive performance can produce substantially different, and in some cases misleading, explanations for the same observations. To systematically identify such cases, we introduce a novel discrepancy-based measure for detecting differences in explanations across models. Building on this measure, we propose a framework for flagging scenarios in which high predictive accuracy masks unreliable explanatory behaviour and identifying scenarios with high explanation consistency. We provide practical recommendations for the effective use of explainable ML in ecological modelling and conservation decision-making. Although illustrated using a marine ecology case study, the proposed framework is broadly applicable across ecological systems and contributes more generally to explainable ML research.

## Methods

We develop a general quantitative framework, inspired by the similarity analysis of explanations by Gwinner et al. (2024) to detect model predictions with high explanation discrepancy across models. This framework enables practitioners to: identify consistent explanations across models; and detect predictions with extreme levels of inconsistency. Both of these tasks are important for reliable decisionmaking as they relate to a form of uncertainty in model explanations, which is distinct to uncertainty in predictions. This section first develops the foundational structure of this framework and then presents a specific implementation of the framework for a popular state-of-the-art explainable ML method. This implementation is utilised in the simulation study presented in Section 3, and alternative implementations are provided in Section S1.3.

### 2.1 An explanation discrepancy framework

We introduce the *explanation discrepancy framework* (EDF) Figure 2) that enables detection of differences of explanations across multiple models. We define an explanation as a real-valued vector *S* ∈ ℝR^*K*^, where each of the *K* ≥ 1 dimensions corresponds to a feature of the data that consists of *N* observations. EDF is based on a distance metric induce by a vector norm ∥ · ∥ on the explanation vector space. Given a set of models 𝕄, let 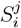 be the explainability vector for the *i*th observation, or prediction, under the *j*th model.

**Figure 2.**
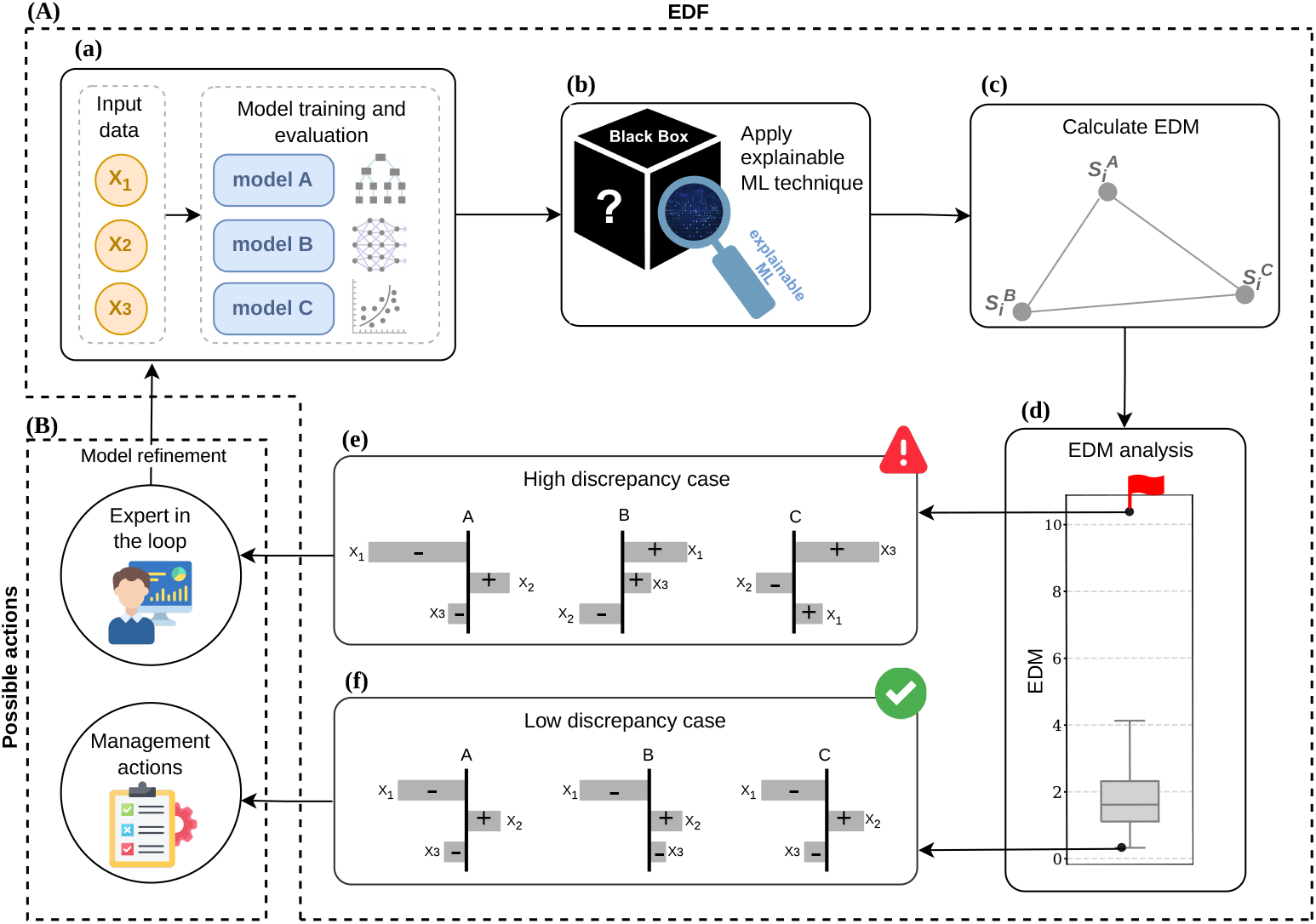
(A) Flow diagram of the explanation discrepancy framework (EDF). (B) Examples of possible actions that could result from the EDF. (a) Multiple ML models (*A, B*, and *C*) are fitted using given input data with features *X*_1_, *X*_2_, and *X*_3_ and they are calibrated to achieve equivalent predictive performance. (b) An explainable ML method is applied to the fitted models. (c) The resulting explainability vectors are then used to compute the explanation discrepancy measure (EDM). (d) Predictions with high EDM are flagged. (e) These observations with high discrepancy in explanations should be directed to domain experts for further investigation whereas, (f) explanation associated with low EDM suggest agreement across models, increasing the robustness of explainability.

We define, for a given observation *i* ∈ {1, 2, …, *N*} and two models *j, k*∈ 𝕄, the relative explanation discrepancy for model *k* relative to model *j*,

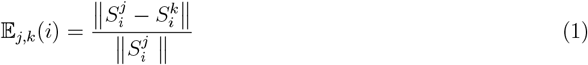

This metric provides insight into the overall inconsistency between a given set of models and follows properties of vector norms 𝔼_*j,k*_(*i*) = 0 if and only if 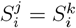. Building on Eq 1 we define the *explanation discrepancy measure* (EDM) as,

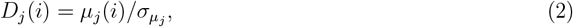

where,

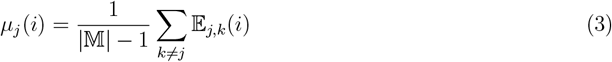

and

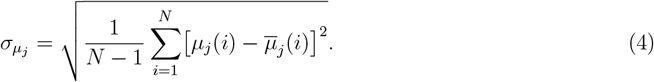

The EDM effectively provides a signal to noise ratio in the relative explanation discrepancy space and we use the EDM to identify scenarios where there is disagreement on explainability for prediction versus the explanations with consensus.

Particularly, we are interested in the predictions with large extreme EDM values (outliers of EDM’s distribution), which indicate strong disagreement in the explanations between models (Figure 2(e)), relative to a particular reference model. Such high discrepancy points indicate instances where the human involvement is crucial and should therefore be directed to domain experts for further investigations. In contrast, lower EDM values indicate higher agreement in the explanations across the models (Figure 2(f)), with respect to a particular reference model, which leads to greater confidence in the models. Therefore, it is equally important to identify where such consistency exists, since that is where an ecologist makes decisions.

As formulated in Eq (1)–(4), our EDM is computed on the full explanation space that includes all input features, some of which may not be of interest. We can restrict our analysis to include only features of interest via projections, leading to a projected EDM. We do not focus on this here, but provide additional details in Section S1.1.

### 2.2 Shapley additive explanations

Some popular explainability methods include SHapley Additive exPlanations (SHAP; Lundberg and Lee, 2017), Local Interpretable Model-agnostic Explanations (LIME; Ribeiro et al., 2016), Individual Conditional Expectation (ICE), Accumulated Local Effects (ALE) plot, Feature Interaction, Feature Importance, and Global Surrogate (Molnar, 2019). In this work, we focus on SHAP, one of the most popular post-hoc model agnostic methods (See Section S1.2 for more details on types of explainable ML methods.) in the current state-of-the-art (El Assari et al., 2023; Rozemberczki et al., 2022), as a specific implementation of our EDF (See Section S1.3 for the mapping of LIME results for the explainability vector.).

Lundberg and Lee (2017) proposed SHAP as a unified framework for interpreting predictions of complex models. This method is inspired by methods to evaluate player contributions in a cooperative game (Shapley, 1953). Given *f* is a flexible ML model and *x* is the set of all input feature values for a particular prediction, the SHAP value denoted by *ϕ*_*i*_(*f, x*), represents the impact of feature *i* on the prediction of *f* for input *x*. When *f*_*x*_(𝕊) = *E*[*f* (*x*) | *x*_𝕊_] is the expected model output for a subset of the input features S which excludes feature *i*, SHAP values are computed by summing over all possible subset 𝕊, with a weighted factor 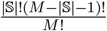 that accounts for impact of each feature being added to the model (Eq 5).

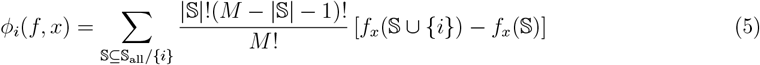

One useful property of SHAP is the local accuracy property expressed as,

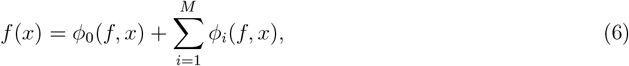

where *ϕ*_0_(*f, x*) = *E*[*f* (*x*)] is the baseline value or the expected model prediction over the training dataset, and *M* is the total number of features in the model. According to this property, SHAP values collectively capture the difference between the prediction *f* (*x*) and the expected model prediction.

The missingness property states that features missing in the input get zero attribution or no effect (*ϕ*_*i*_(*f, x*) = 0), while the property of consistency indicates that if a model changes such that an input features’s contribution increases, then regardless of the other inputs, the importance attributed to that feature should not decrease. Moreover, large positive or negative SHAP values convey a significant effect on the individual prediction by the given feature in the model, whereas the sign of the SHAP value tells us the direction of that effect, whether it is positive or negative.

In this work, SHAP values are computed at the prediction level from the set of models *M* and in our SHAP analysis, we focus mainly on different summarisation methods such as waterfall plot, beeswarm plot (Figure 3), and mean SHAP. Waterfall plot, as a local explanation method, visualises how each feature has contributed to a specific local prediction, from the baseline or average prediction. For example, Figure 3(a) illustrates a single prediction of abalone age, modelled by a ML model using five input features (e.g., height, length, etc). Thus, for each individual observation, we can observe how the prediction is being made by the features involved in the model.

**Figure 3.**
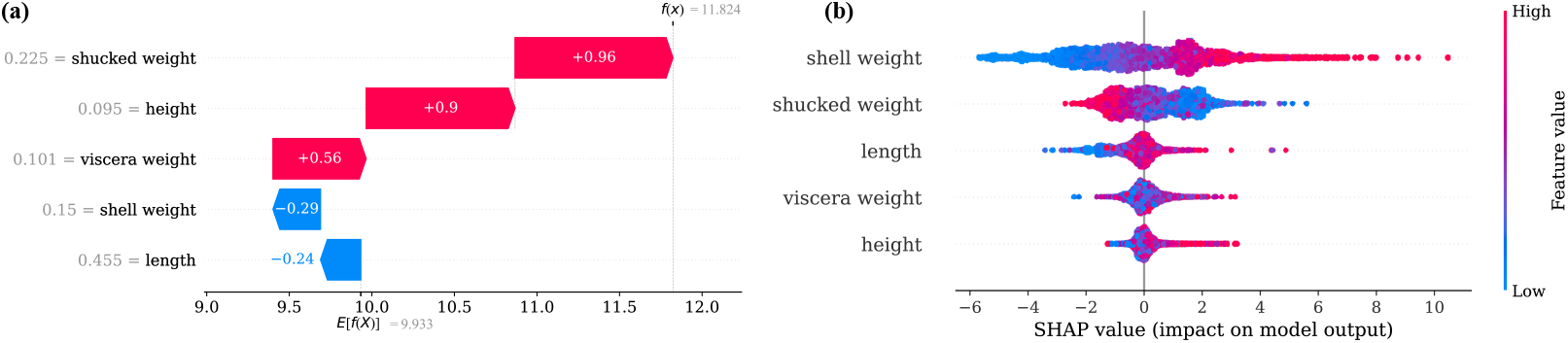
Example SHAP (a) waterfall plot and (b) beeswarm plot, using the Abalone dataset obtained from https://archive.ics.uci.edu/dataset/1/abalone. The age of abalone is predicted by a boosted regression trees (BRT), using five features of the related physical measurements of Abalone. (a) explains a single prediction of abalone age using the features in the model. Features are listed in the y-axis. The x-axis represents the SHAP values and the sign indicates the effect on the prediction by features, whether it is positive (increasing) or negative (decreasing). (b) visualises SHAP values of all observations (4177) for each feature (y-axis), where the x-axis represents SHAP value. Each dot corresponds to a single prediction. The colour scale indicates the value of the the feature – high (red) or low (blue).

The beeswarm plot (Figure 3(b)) visualises the SHAP values across all observations, while mean SHAP provides the averaged absolute SHAP values for each feature across all those observations. As a bar plot, the mean SHAP depicts which features are most important to make the predictions by the model. Features with larger positive or negative SHAP values result in larger mean SHAP, indicating that those features have a substantive impact on the prediction given by the model, making the mean SHAP to be a representative of feature importance. In light of this, SHAP is not only a local explainability method, but is also capable of providing global explanations (Raschka et al., 2020).

## 3 Simulation study

We demonstrate the utility of EDF in a setting where the ground truth is known. To establish a ground truth, we generate synthetic data that enables us to explore the implications of explainability sensitivity for the determination of environmental drivers on population dynamics. We fit four models, including three ML based models and a statistical approach, to these synthetic data to evaluate reliability and consistency of explanations against the known associations in the data. We also compare explainability results with classical statistical inferences.

### 3.1 Synthetic data generation

We consider synthetic data that results from a stochastic mechanistic model with known ground truth behaviour. This ground truth is important to evaluate the quality of explanations, as this is not possible with real data, especially for complex ecosystems such as coral reefs. We generate semi-realistic synthetic coral cover data using *Synthos*, an R software introduced by the Australian Institute of Marine Science (AIMS) (Logan and Vercelloni, 2025). It simulates three mechanistic disturbances: cyclones, heat stress (bleaching), and other more-localized disturbances in which their intensities are controlled. We develop on a cyclone-dominated scenario using the weighting disturbance system incorporated in *Synthos* that controls the relative influence of the disturbances on the spatio-temporal variability of coral cover. Coral cover is modelled in *Synthos* as a cumulative effect of a baseline coral cover value that represents the cover before sampling, weighted disturbances, and annual growth of coral. Thus, using *Synthos*, we generate yearly values of coral cover for 30 years across a synthetic predictive domain and we select data for 49 reefs (with 3 sites per each reef) including coral cover and disturbance values for model training and testing purposes.

Figure 4(c) represents an example coral cover trajectory from 147 sites following a simulation of the protocols implemented by AIMS’ long term monitoring program. In this way, we have some data in which we know the effect of disturbances on coral cover over time (Figure 4(b),(c)). Section S2.1 provides further details relevant to this simulation, including the resulting trajectories of mean hard coral cover (MHCC; hard coral cover averaged across transects) for a depth of 10m, which is one of the depths in the simulation. For our modelling framework (Section 3.2), we utilise this data from all reefs. In this application, our features used as input to the models to predict the MHCC at site level are the covariates shown in Table 1 and they are standardised when the models are built.

**Table 1.**
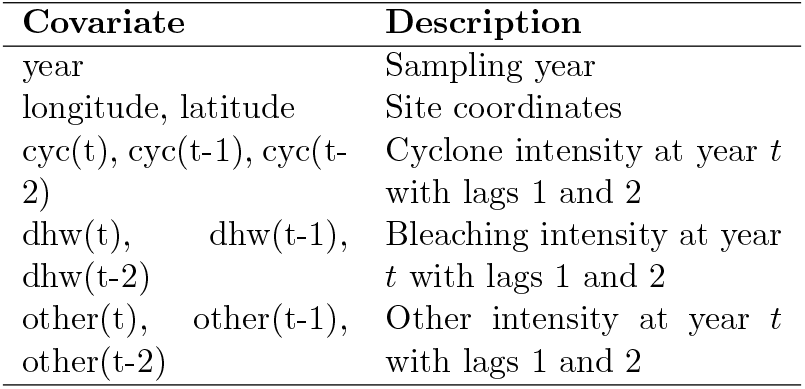
Description of covariates considered under this study to model the mean hard coral cover (MHCC). The disturbance intensities refer to the relative intensities of disturbances before weighting.

| Covariate | Description |
| --- | --- |
| year | Sampling year |
| longitude, latitude | Site coordinates |
| cyc(t), cyc(t-1), cyc(t-2) | Cyclone intensity at year $t$ with lags 1 and 2 |
| dhw(t), dhw(t-1), dhw(t-2) | Bleaching intensity at year $t$ with lags 1 and 2 |
| other(t), other(t-1), other(t-2) | Other intensity at year $t$ with lags 1 and 2 |

**Figure 4.**
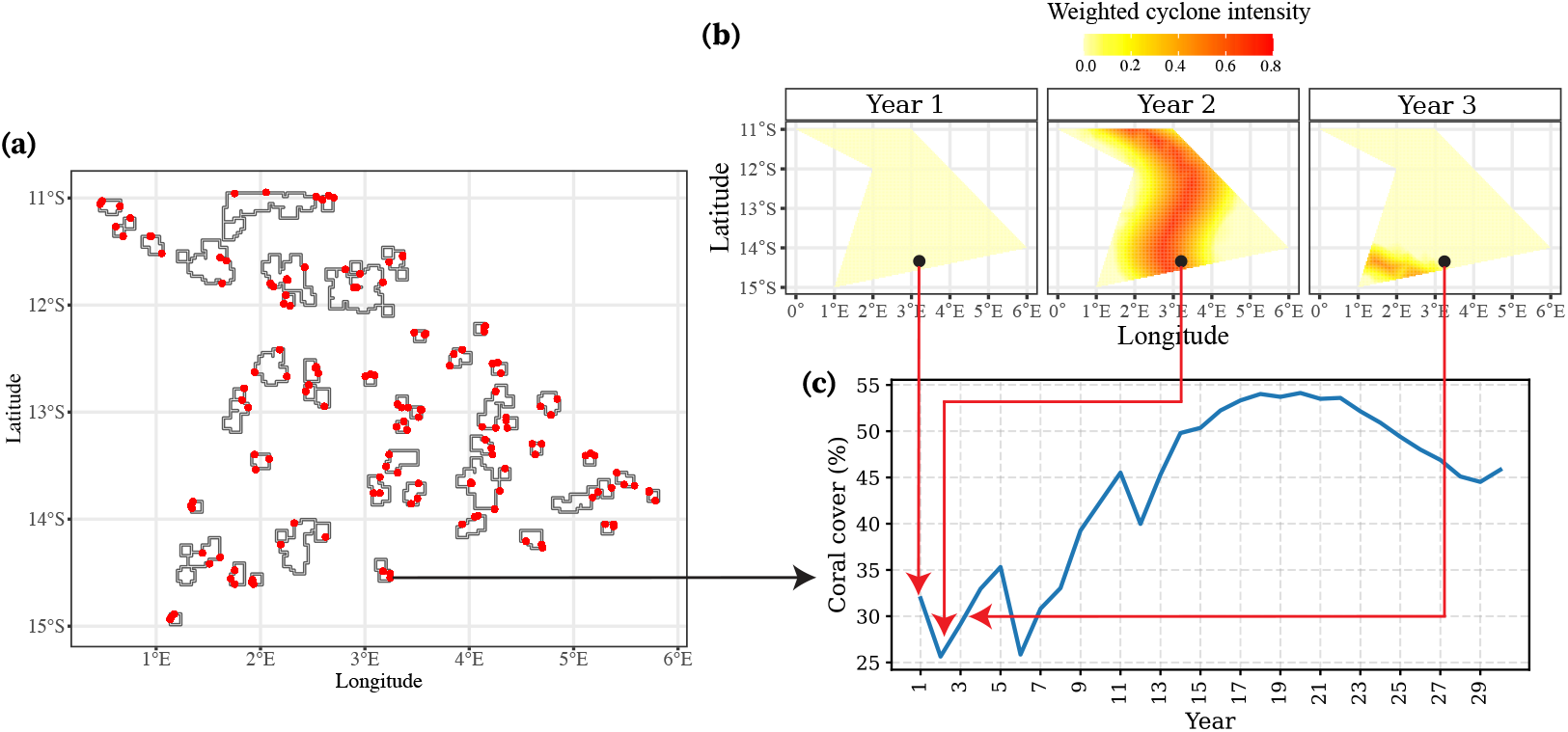
Schematic of synthetic data generated from *Synthos*. (a) represents the locations of the synthetic reefs in the defined spatial domain, where the irregular shaped polygons represent the reefs and the monitoring locations are indicated from red dots. In total, there are 49 reefs, with 3 sites for each reef. (b) shows heatmap of one of the mechanistic disturbances after applying the corresponding weight (weighted cyclone) generated for three years and the black dot represents one of the sites. (c) provides the generated coral cover over time for that particular site (see Figure S9 for the generated coral cover over time for all sites). For example, the arrows correspond to the first second and third year in this time series, and the coral cover decline in year 2 is associated with some known mechanistic disturbance effect.

### 3.2 Modelling frameworks

To demonstrate our EDF, we fit a selection of models to the synthetic coral data (Section 3.1), using different ML and statistical methods. These ML methods include: random forest (RF), boosted regression tree (BRT), and artificial neural network (ANN). The statistical approach we use is the generalised additive model (GAM). In this section, the selected methods are described, followed by model construction and evaluation.

#### 3.2.1 Generalised additive models

Generalised additive models are a flexible class of statistical models of the form,

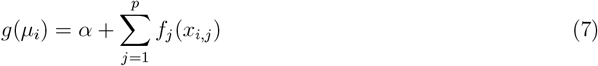

where *x*_*i*,1_, *x*_*i*,2_, …, *x*_*i,p*_ are *p* features and *µ*_*i*_ = *E*[*Y*_*i*_] is the mean response, *g*(·) is the link function that maps *µ*_*i*_ to the real line, *f*_1_(), *f*_2_(), …, *f*_*p*_() are flexible smooth functions (Simpson, 2018) and *α* denotes the intercept. One of the major strengths of GAM is its ability to deal with non-linear relationships between response and features, while retaining its interpretability (Doohan et al., 2026). Since GAMs are inherently interpretable via statistical inference, have been a standard tool in ecosystem modelling for decades (Guisan et al., 2002; Lai et al., 2024; Singh et al., 2024).

#### 3.2.2 Random forests

RF is a decision tree-based modelling approach in ML and is considered to be one of the most precisely performing models in ecological studies (Mi et al., 2017; Shabani et al., 2016). RF combines multiple regression trees, each of which is trained and validated on a random subset of the data (Breiman, 2001a; Algorithm S1). RF predictions are based on averages across individual tree predictions (Schonlau and Zou, 2020; James et al., 2013). While RF is very flexible and can capture complex non-linear relationships (Canning and Waltham, 2021), the algorithmic nature of the method lacks explicit interpretability (Simon et al., 2023).

#### 3.2.3 Boosted regression trees

BRT is another widely used ML algorithm in ecological modelling. It is a powerful tool that involves many decision trees, where each of these trees are combined in a forward and stepwise fashion (Canning and Waltham, 2021). BRT improves the predictive performance of the final model in an iterative way (Algorithm S2) that enables learning from the errors by predicting the residuals of the previous tree at each subsequent tree (Pittman and Brown, 2011). While BRTs facilitate significant flexibility to deal with complex non-linear relationships, similar to RF, they are considered as “black-boxes” as they are not easily interpretable (Welchowski et al., 2022).

#### 3.2.4 Artificial neural networks

ANN is a non-linear modelling technique inspired by biological neural structure of the human brain, consisting of interconnected layers of “artificial neurons” (Ghosh, 2026, Figure S10). In this work, we consider a multilayer perceptron (MLP; Figure S11), which is one of the most frequently used ANN architectures for a wide variety of problems including ecological modelling (Park and Lek, 2016). Managing large volume of input data, ability to represent various forms of complexities in prediction tasks, and fast learning capability are some of the popular advantages of MLP (Hu et al., 2024). However, a common criticism of MLPs is that interpreting the inner structure of the function they learn is challenging, leading them being defined as “black-box” methods (Swingler, 2014; Altuhaifa and Al Tuhaifa, 2025).

### 3.3 Model construction and evaluation

We split the simulated data explained in Section 3.1 for our model training. As indicated in Table 1, we consider two disturbance lags to model coral cover and therefore, associated data from year 3 are incorporated into the analysis. Thus, we select a random subset (70%) of observations from year 3 to 29 for training, and use the remaining 30% and the first forecast year (year 30) for testing. Considering random subsets of observations for model training and testing allows gaps in both space and time, avoiding loss of important information (signals) on disturbances which is useful for model training. In addition, this test strategy facilitates a training set that covers a range of disturbances that helps predict coral cover under individual disturbances or combinations of disturbances, allowing for a fairly representative sample for model training and testing.

GAMs are built using the pyGAM package in Python (Servén and Brummitt, 2018), including spline-based smooth functions and using Normal distribution with identity link function. We perform randomised grid search over a range of smoothing parameter (*λ*) values that control the strength of the regularization penalty on each term or the wiggliness of the fitted functions, and automatically tune the model using generalized cross-validation (GCV; Wood, 2017). The corresponding equation of the GAM that fitted our data well is shown in Eq 8, where *µ* represents the MHCC at site level for a given year, *α* is the intercept, *s*(.) and *te*(.) denote spline smooths and tensor product smooth interactions, respectively. In order to compare with the ML methods, the same set of features associated with the GAM are utilised as features to the considered ML methods. All three ML models are tuned using random search with 5-fold cross-validation on training data, involving a range of hyperparameter combinations (Section S2.3). To evaluate the models in terms of their predictive performance, we use a range of standard metrics (Eq S6–S9) including, mean absolute percentage error (MAPE), root mean square error (RMSE), mean absolute error (MAE), and coefficient of determination (*R*^2^).

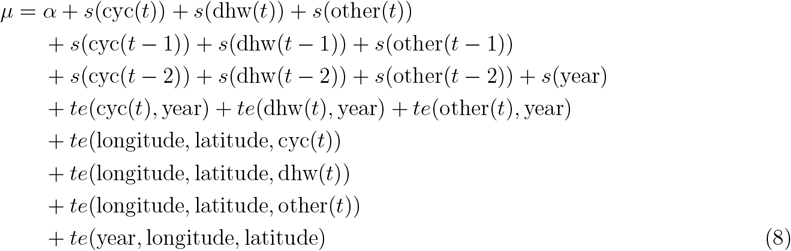

## 4 Results

In this section, we present the main results of our analysis. To ensure valid comparisons and conclusions from EDF, we ensure that all models perform equally well in terms of predictive performance under the standard metrics described in Section 3.3 (see Section S3.1). We then present results based on SHAP, and use our novel EDF to identify observations of extreme disagreement between the ML explanations and consistencies as well. Finally, we evaluate the utility of our EDF using three scenarios available within the *Synthos* dataset.

### 4.1 SHAP reveals consistent global explanation patterns across models

As a global explanation, the beeswarm plots of four models for the within-sample analysis (Figure 5), convey the overall trend the models use to make predictions in general. Each point in the beeswarm plot represents a single site in a specific year. Therefore, this plot can be used to explore the nature of the relationships between different features used to model the MHCC.

**Figure 5.**
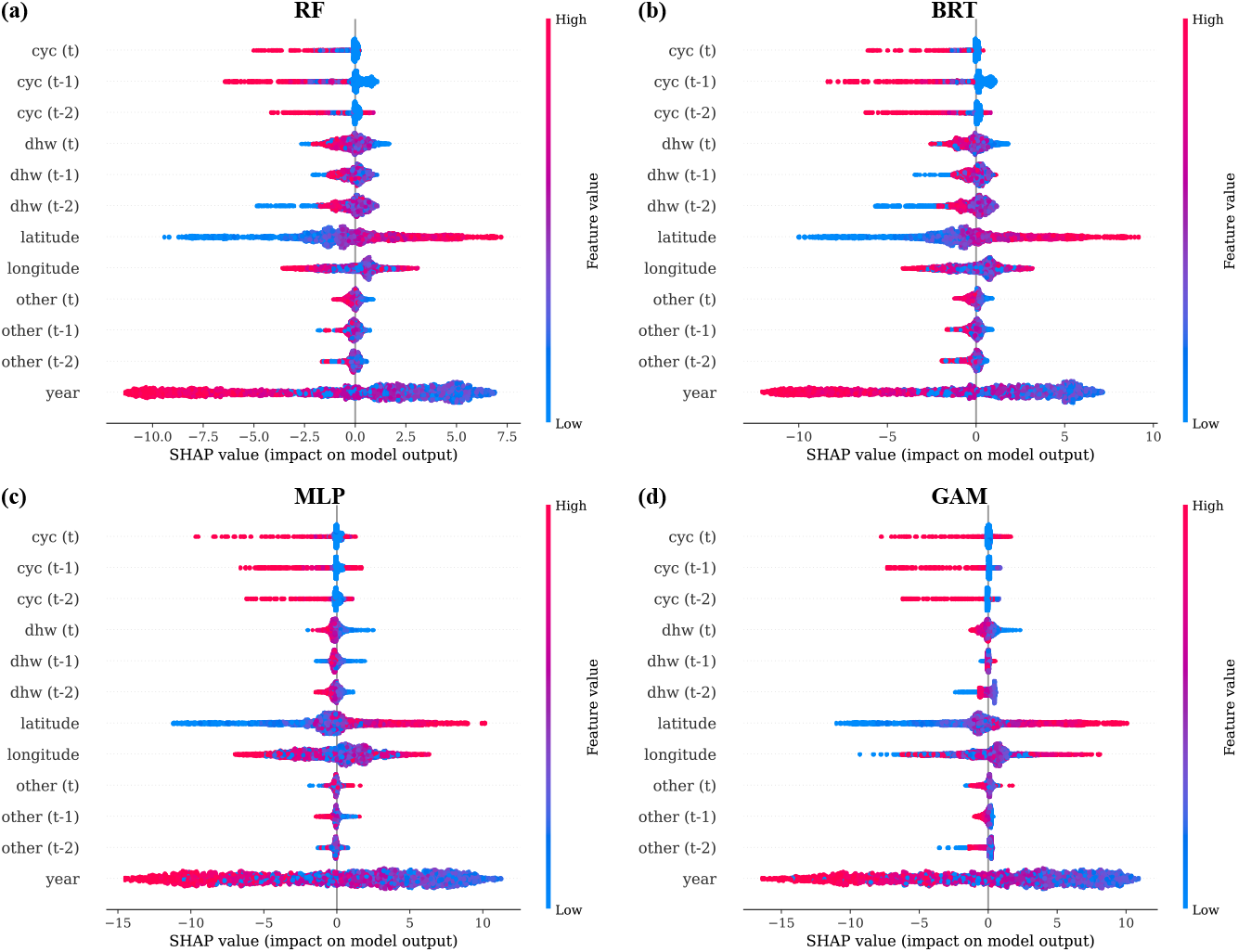
Beeswarm plot for (a) RF (b) BRT (c) MLP, and (d) GAM for the within-sample (training) data. The y-axis shows the features in the model and they are ordered alphabetically, for direct comparison, rather than the standard ordering by importance. The x-axis indicates SHAP value. Each dot represents a single site of a reef for a particular year. The absolute higher SHAP value indicates a higher effect on the model output or the prediction. The sign indicates the effect on the prediction, whether it is positive (increasing) or negative (decreasing). The magnitude of the SHAP values, expressed using a colour scale from red (high) to blue (low), is an indicator of how strong the effect of a feature is on the individual prediction of MHCC.

Overall, global explanations are largely consistent across the models within-sample (Figure 5), which is also true for the out-of-sample (Figure S14). In terms of the feature importance in the models (Figure S15), the three most influential features for prediction of the MHCC (year, latitude, and longitude) are consistent across the four models. In particular, according to Figure 5, when compared with SHAP patterns of current and lagged dhw and other disturbances, cyclone exposure and associated lags reveal that the general trend or the known ground truth of disturbance weights has been captured by the models. Our synthetic data set exhibits a realistic steady decline in MHCC over time (Figure S9). A similar pattern is identified with the global SHAP analysis across all models with higher values of the year feature tending to lead to negative SHAP values. In addition, significant contributions from latitude and longitude are visible through their higher SHAP values in both directions across the four models, representing the crucial role played by the spatial component in *Synthos*. Altogether, this demonstrates an alignment of global SHAP explanations with our simulator, reflecting the cyclone dominated scenario as described in Section 3.1.

### 4.2 EDM indicates explanation uncertainty

Despite the globally consistent SHAP patterns described in Section 4.1, we found that there is a possibility of different models having significant discrepancies in terms of local explanations. Using the SHAP implementation of the EDM distribution across all observations (Figure 6(a)), we identify three representative scenarios: a high EDM common outlier across all reference models (point *E*_1_ in Figure 6(a)), and two examples of observations with consistently low EDM across all reference models (*E*_2_ and *E*_3_ in Figure 6(a)). For all these three cases, we have a clear ground truth effect obtained through the *Synthos* simulation. As a result, we can evaluate the relationship between EDM and consistency with this ground truth. Each case is visualised with a combination of outputs, including time series plot of observed and predicted values from the four models (to show where the respective point occurred in time), relevant heatmaps (with 2 time lags) of weighted cyclone intensity, and corresponding waterfall plots from the four models. The heatmaps help understand where the points occur in space and the magnitude of actual disturbance effects on them. We present this combination of plots as a recommended layout for data point diagnostics to analyse explanations.

**Figure 6.**
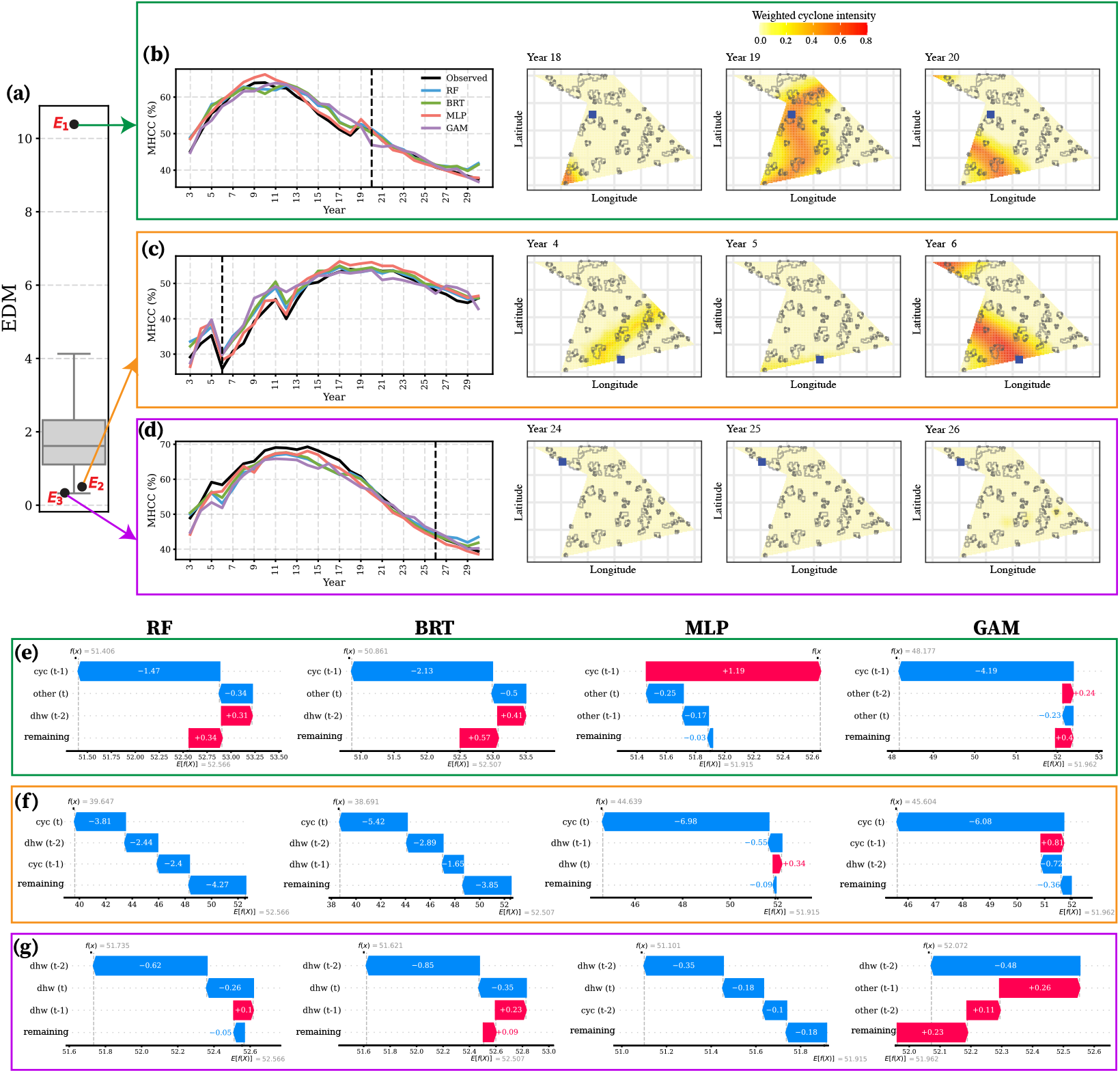
Three representative examples of EDM analysis implemented with SHAP. (a) Box plot of EDM (See Figure S16 for the reference-based distributions of EDM implemented with SHAP). *E*_1_ (site 1 of reef 35, year 20) represents a high EDM outlier, *E*_2_ (site 1 of reef 25, year 6) represents a clear cyclone event with low EDM value, and *E*_3_ (site 2 of reef 44, year 26) represents a data point with low EDM value. (b)-(d) The left panel illustrates the time series plot of observed and predicted values by the four models at the respective site. The vertical dashed line indicates the particular data point (*E*_1_-*E*_3_). The right panel shows heatmaps of weighted cyclone intensity with two time lags, where the grey coloured dots indicate the sites of each reef (irregular polygons) and, the focal site is highlighted with a blue coloured square. (e)-(g) Corresponding waterfall plots from RF, BRT, MLP, and GAM models for *E*_1_, *E*_2_, and *E*_3_, respectively. These plots visualise the individual contribution of features involved in the model to the final predicted mean hard coral cover (MHCC), from the expected model output *E*[*f* (*x*)] over the training dataset. The y-axis shows the features in the model, whereas the x-axis indicates SHAP value. The horizontal bars convey the direction of the contribution of the feature, whether it is positive (red) or negative (blue). We focus on the top feature of these waterfall plots once longitude, latitude, and year are removed. The corresponding full waterfall plots and other two disturbances are shown in Section S3.3.

In Figure 6(a), *E*_1_ corresponds to a higher EDM value and is identified as an outlier in the EDM distribution. As shown in the heatmap of Figure 6(b), the ground truth associated with *E*_1_ exhibits that this particular site has been affected from cyclone in the previous year (year 19). According to the local SHAP explanation for *E*_1_ (Figure 6(e)), RF, BRT, and GAM indicate that lag 1 of cyclone contributes negatively to predict the MHCC, whereas MLP suggests the opposite which is inconsistent with the simulator.

On the other hand, *E*_2_ in Figure 6(a) depicts a data point with a clear cyclone event and low EDM value. The significant drop in MHCC in year 6 that can be seen in the time series plot associated with *E*_2_ (Figure 6(c)) is caused by the cyclone at that year according to the simulator. In terms of the local SHAP explanations of *E*_2_ (Figure 6(f)), regardless of the model type, the most dominant feature, cyc(t) shows a negative effect on the predicted MHCC indicating that cyclone at year 6 has been a major factor to the significant decline in coral cover. That is not only consistent across all models but also aligns with our simulator.

Moreover, *E*_3_ represents a data point with a low SHAP discrepancy. It is evident from the corresponding heatmaps, the other two disturbances (dhw and other) are the influential factors for decline in MHCC (see Figure S19(f)–(h) for comparable disturbances) for the underlying year and its two lags. According to the local explanations (Figure 6(g)), cyclone or its lags do not emerge as dominant features, suggesting an alignment of the model explanations with the known ground truth from our simulator.

Thus, these three scenarios suggest that when the EDM is small, the models behave similarly in terms of their local explanations and are also consistent with our simulator, while data points with high EDM exhibit noticeable disagreements in the explanations across the models, deviating from the known relationships regardless of equivalent predictions (Figure S20 further emphasises that SHAP discrepancy is uncorrelated with prediction discrepancy) between the models.

## 5 Discussion

In this work, we introduce a new framework that facilitates a systematic approach to dealing with the sensitivity of explanations to the model with some reliability in ML models. Through the use of simulated coral reef data from *Synthos*, we investigate the potential of EDF to estimate the explainability and sensibility of ML methods across multiple models. Our analysis is inspired by modelling efforts of coral reefs, which are some of the most threatened ecosystems globally (Hughes et al., 2017, 2018; Warne et al., 2022, 2025; Ortiz et al., 2018; De’Ath et al., 2012; Bozec et al., 2025). There has been substantial growth in the use of ML methods in the study of coral reefs (McClanahan, 2025; Jouffray et al., 2015; Couce et al., 2013; Mellin et al., 2019; Cheung et al., 2025). Due to the fact that management and restoration actions act on the underline drivers (Darling et al., 2019; Muñiz-Castillo et al., 2024; Facon et al., 2016), predicting changes in coral abundance does not provide sufficient information to support conservation efforts if the drivers of coral decline are not identified.

Furthermore, given the massive costs associated with the management of coral reefs, including coral restoration and adaptation (Bayraktarov et al., 2016; Darling et al., 2019; Anthony et al., 2020; McLeod et al., 2022; Bay et al., 2023), quantification of drivers needs to also be associated with uncertainty, which ML alone cannot achieve. Taken together, coral reefs can be considered as an ideal ecosystem for the implementation of explainable ML methods and the proposed framework.

It is apparent that there is a possibility of different models having significant discrepancies in their explanations corresponding to the same local observations (*E*_1_ in Figure 6). In particular, when the different models are linked to different conclusions of how the prediction is being made for the same observation, it raises the question of how we should rely on these ML methods in an ecological context in the absence of a measure of explanation uncertainty.

Our EDF enables identification of points of disagreement between ML models, and more broadly between any two models with same features, where an expert should be consulted for further analysis on that. Our key message based on our results is to focus on the predictions where the EDM is large, as it indicates a significant disagreement in the explanations across different models. If a particular prediction is flagged as a common outlier for more than one reference model (similar to the scenario of *E*_1_ in Figure 6), it can be considered as a consistent or strong outlier of explanation discrepancy, where further exploration on the scenario is crucial. This study also highlights that even if all models have similar fit and accuracy (Section S3.1), we can expect substantial differences in explanations locally, which is a great motivation to focus on the EDF. When the EDM is small (i.e., *E*_2_ and *E*_3_ in Figure 6), the models tend to agree more in terms of their explanations aligning with the ground truth. Thus, if all the models describe the same trend, it helps build confidence in the ML models and their explanations as a reliable characterisation of the ecological processes underlying the data. Therefore, unlike using a single ML model, focusing on multiple models and comparing their explanations seem to be a promising approach for a reliable use of explainable ML.

While we demonstrated an implementation of our framework using SHAP and its applicability to LIME (Ribeiro et al., 2016) in Section S1.3, our approach is generally applicable to any local model-agnostic method that assigns numeric values to features. Successful implementation of EDF for multiple explainability methods will allow evaluating the model consensus across multiple explainers, in addition to within-explainer comparisons. Furthermore, examining the potential impact of different environmental intervention strategies through the use of EDF will largely benefit conservation of ecosystems by providing more ecologically informed information to different stakeholders, including reef managers.

The current EDF does not incorporate any uncertainty quantification to deal with the potential uncertainty associated with discrepancy in explanations. To address this gap, future work can also include extending this framework accounting for uncertainty in the model explanations in the Bayesian setting. In addition, while the lower and upper tails of the distribution of EDM represent consistent and inconsistent explanations, respectively, there is currently no systematic approach or threshold for the classification of explanations into different regions to represent their level of consistency.

Given our findings, we caution against the standard practice in ecology of using explainable ML for a single ML model. Limiting to one model that has shown excellent predictive performance is not necessarily the ideal option. Instead, exploring multiple methods and their explainability results is recommended as an advisable way to identify inconsistent local and global explanations obtained through different models. Particularly, as equivalent predictability need not imply equivalent explainability, there is a risk associated with decision-making based on the conclusions derived from an explanation obtained from a single ML model. In this setting, there is no principled approach to determine if the patterns identified by the model are plausible patterns in terms of real world. To overcome this, we recommend to use more than a single ML model with the EDM to identify reefs or regions that require further attention before implementing any conservation action.

The findings of this study highlight the extent to which explainable ML methods provide real-world ecological insights that can be utilised for realistic management decisions. More importantly, this novel framework is not limited to applications related to the marine space but can also be generalised to different ecosystems and different research disciplines. It also leads to the effective use of ML methods with confidence, given the sensitivity of the explanations provided by explainable ML methods. To date in the ecology literature, there is no principled framework to evaluate the differences in explanations across multiple models and therefore, bridging this gap, our EDF contributes to the existing body of knowledge of explainable ML as a valuable addition.

## Acknowledgements

AD is supported by a Research Training Program Scholarship. DJW is supported by an Australian Research Council Discovery Early Career Researcher Award (DE250100396).

