## Supplementary Appendix for "An explainable machine learning consensus framework for robust estimations of environmental effects on population dynamics"

### 1 Additional information for Section 2

#### 1.1 Projected explanation discrepancy measure (Projected EDM)

In *Synthos*, coral cover dynamics heavily depend on space and time. Therefore, year, longitude, and latitude are considered as strong factors affecting coral cover, regardless of whether there is a disturbance

or not. While these factors should appear as features in the models as they are strong predictors, we may not need to focus on them in model explanations. This motivated calculation of the discrepancy in explanations, removing particular dimensions that are not of interest. The resulting discrepancy is defined as "Projected EDM". It can be used to identify differences in model explanations for ecologically interested features such as disturbances. Further results (Figure 1) suggest that the Projected EDM analysis implemented with SHAP (Projected SHAP discrepancy) does not show any strong correlation with the EDM analysis implemented with SHAP (SHAP discrepancy) that involves the full SHAP vector.

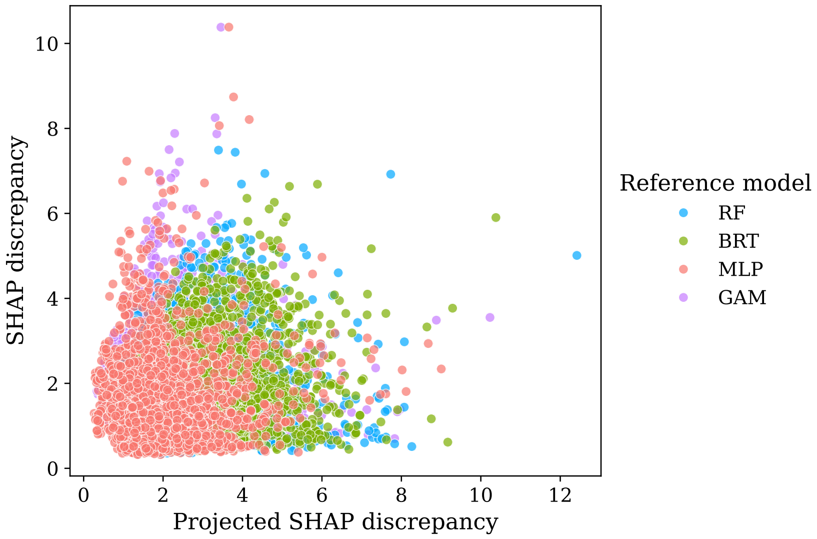

Figure 1: Scatterplot of SHAP discrepancy versus projected SHAP discrepancy. SHAP discrepancy refers to the full SHAP discrepancy which incorporates all features in the model, while projected SHAP discrepancy is calculated based on the lower dimensional SHAP vector after excluding specific features of no interest in the explanation.

#### 1.2 Explainable machine learning methods

Explainable ML techniques are broadly divided into three categories as intrinsic or post-hoc, model-specific or model-agnostic, and global or local. The behaviour of the entire model is conveyed by global explainability while local explainability relates to explaining individual predictions (El Assari et al., 2023). Post-hoc (i.e. after model fitting) methods analyse the behaviour of fitted complex models, explaining variable importance for predictions and relationships between the response and predictor variables. Some post-hoc methods are applicable to any complex model, making it model-agnostic, meaning that they can be used across different models, while some of them are model-specific or can only be used for a particular model. As these categories are not mutually exclusive, they can be used jointly to explain the same model for different purposes (Ryo et al., 2021).

#### 1.3 Mapping LIME output for the explanation discrepancy measure (EDM)

One of the most important breakthroughs in explainable ML includes the post-hoc interpretation method, Local Interpretable Model-agnostic Explanations (LIME), a novel explanation technique proposed by Ribeiro et al. (2016). LIME is an algorithm that can provide interpretable and faithful explanations about the predictions provided by any classifier, by approximating it locally with an interpretable model. LIME aims to provide explanations on how the fitted complex model reaches a prediction for a given instance by fitting a “local surrogate” or a simple model such as a logistic regression or decision tree. Thus, by approximating the behaviour of the complex model, LIME provides a better understanding about how the complex global model arrives at local predictions.

When $x$ is the original representation of a targeted instance that is being explained, the explanation is defined as a simple model $g$ (i.e., Eq 2), where $g$ belongs to a class of potential interpretable models $G$.

$$\xi(x) = \arg\min_{g \in G} \mathcal{L}(f, g, \pi_x) + \Omega(g) \quad (1)$$

$$g(x_i) = \beta_0 + \sum_{j=1}^{p} \beta_{i,j}(x_{i,j}) \quad (2)$$

The term $\Omega(g)$ (Eq 1) is a measure of complexity of the explanation $g$, represented as the number of parameters. To learn the local behaviour of complex model $f$, samples are drawn around $x$ by purturbing feature values. Given the sample in the original representation and perturbed sample are $z$ and $z'$ respectively, $f(z)$ is used as a label for the explanation model. $Z$ represents the dataset of perturbed samples with the associated labels. $\pi_x(z)$ is a proximity measure between an instance $z$ to $x$, representing a locality around $x$. $\mathcal{L}(f, g, \pi_x)$ (Eq 3) is considered to be a measure of unfaithfulness of $g$ in approximating $f$ in the defined locality $\pi_x(z)$ . As given in Eq 1, in order to get an explanation $\xi(x)$, $\mathcal{L}(f, g, \pi_x)$ is minimised while retaining $\Omega(g)$ low enough so that it can be interpretable by humans (Ribeiro et al., 2016).

$$\mathcal{L}(f, g, \pi_x) = \sum_{z, z' \in \mathbb{Z}} \pi_x(z) \left(f(z) - g(z')\right)^2 \quad (3)$$

In order to map LIME output for the EDM, for a given observation $i$, explanation vector $S_i^j$ is the resulting set of coefficients ($\beta_{i,j}$) for $j$ features from the simple surrogate model $g$ (Eq 2).

### 2 Additional information for Section 3

#### 2.1 Synthetic data generation

Using *Synthos*, we generate synthetic hard coral data which is similar to those obtained from the Australian Institute of Marine Science's long-term monitoring program (LTMP). Considering the point-based method (Hill and Wilkinson, 2004), our sampling design includes 49 reefs, 3 sites per reef, 5 transects per site, 100 photo frames per transect, 50 points per frame, and 2 depths. Figures 3–5 illustrate the spatial and temporal maps of the generated disturbances. In this work, we focus on a cyclone dominated scenario with relative disturbance weights of 0.8, 0.1 and 0.1 for cyclone, bleaching and other disturbances, respectively. Figures 6–8 show corresponding maps of spatial and temporal distribution of actual influence of disturbances once these weights are applied to the generated disturbances. Additionally, annual growth values are set to 3% for hard coral and 3% for soft coral.

Using the fixed survey option (Bray et al., 2025), we monitor fixed locations for 30 years, as we want to simulate the maximum amount of data available from the LTMP. All of these configuration values can be changed as needed due to flexibility of *Synthos*. Once the hard coral cover values are generated, site level mean hard coral cover (MHCC) was calculated by averaging the transect level coral cover. For this study, we use data from all 149 sites and one of the depths (10m) and, Figure 9 shows the trajectories of MHCC at these sites for the 10m depth, which is one of the two depths.

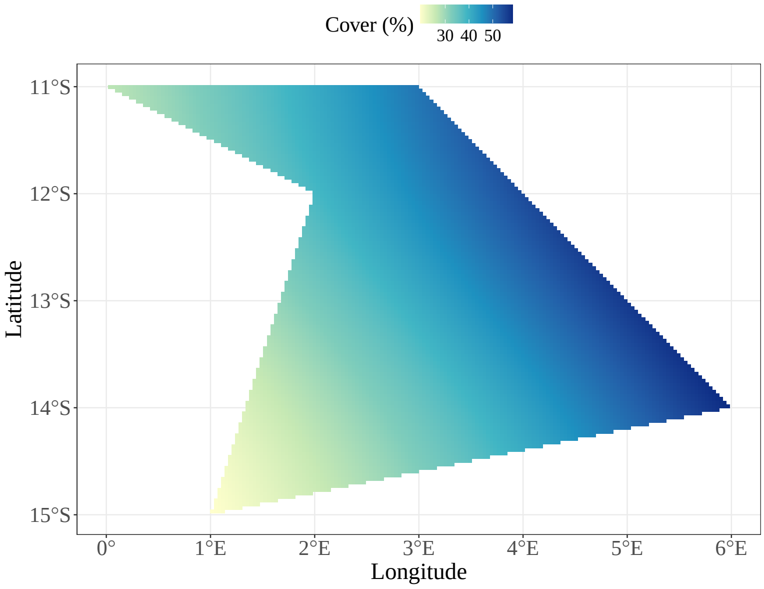

Figure 2: Baseline values of hard coral cover with a west-east spatial gradient over the spatial domain.

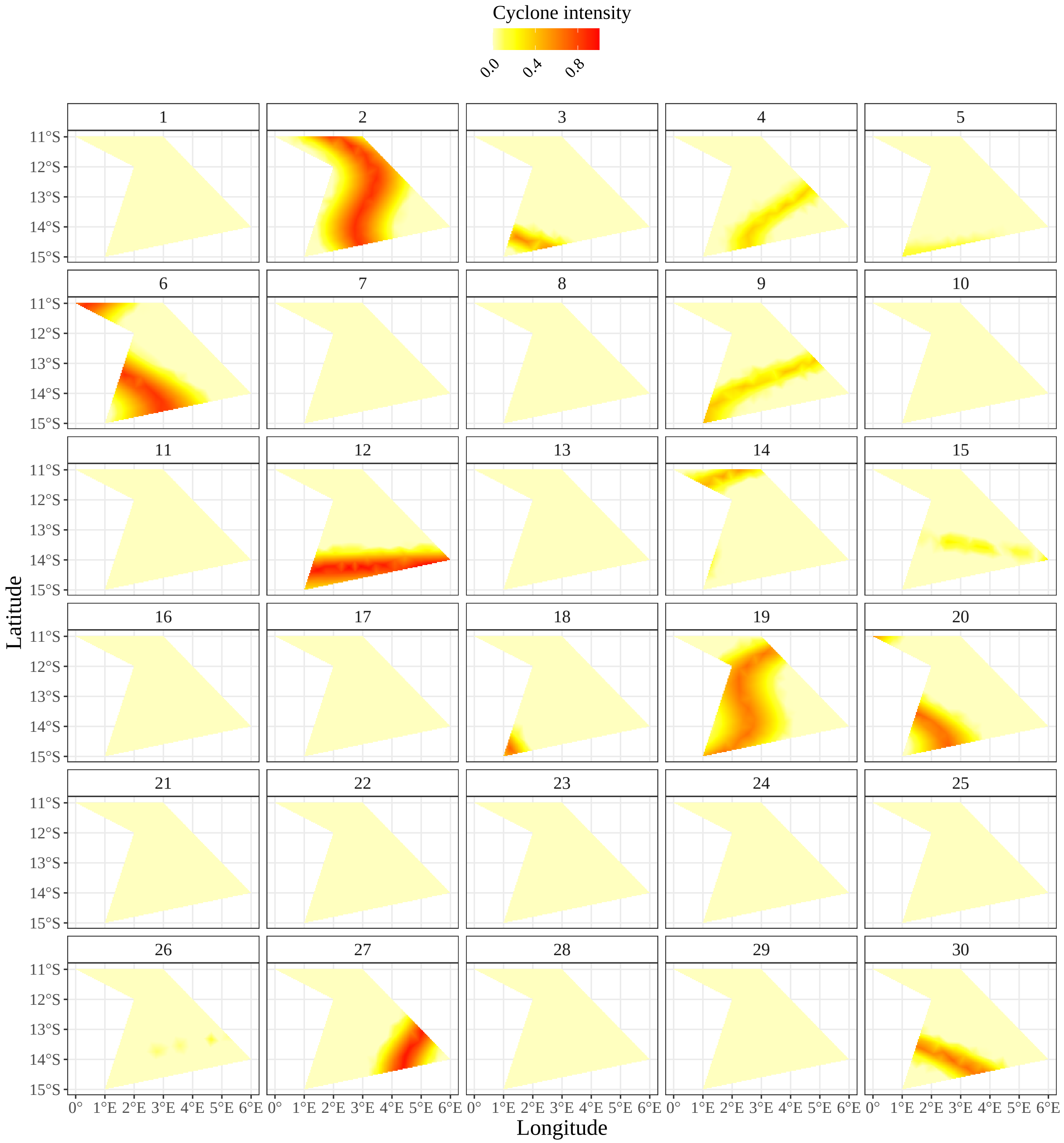

Figure 3: Simulated relative intensity of cyclone in space and time for the 30 years.

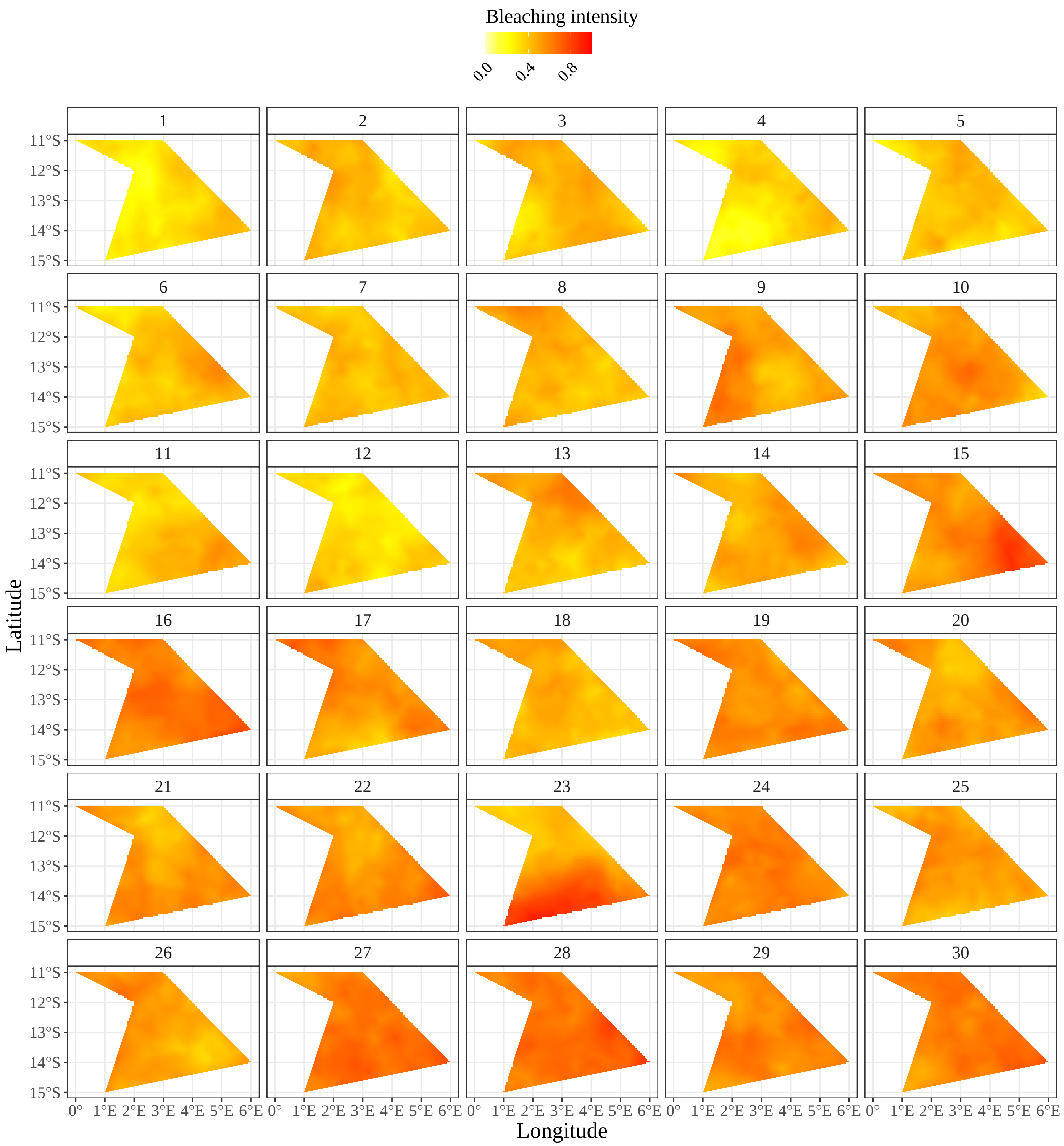

Figure 4: relative intensity of heat stress or bleaching in space and time for the 30 years.

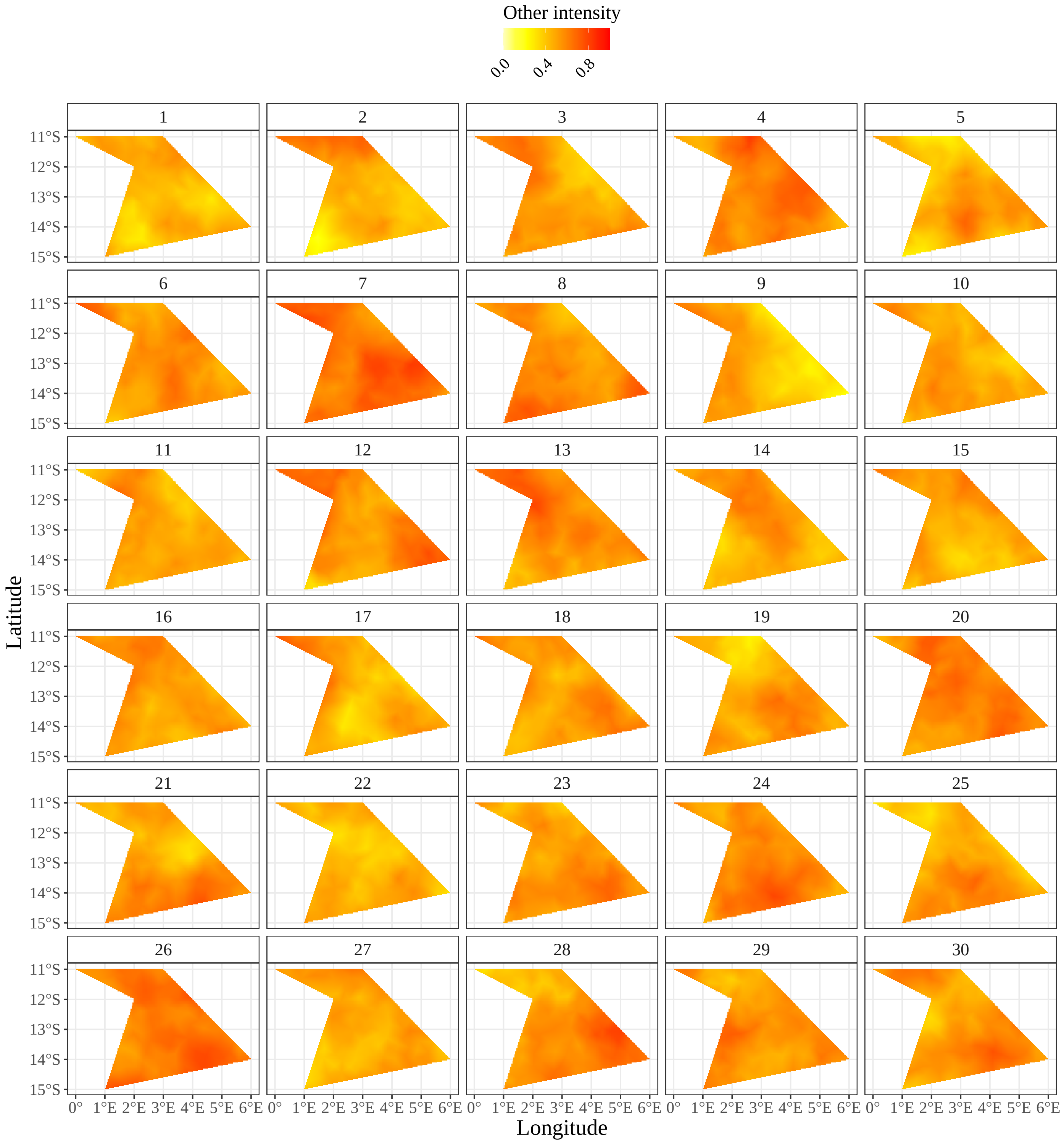

Figure 5: Simulated relative intensity of other disturbances in space and time for the 30 years.

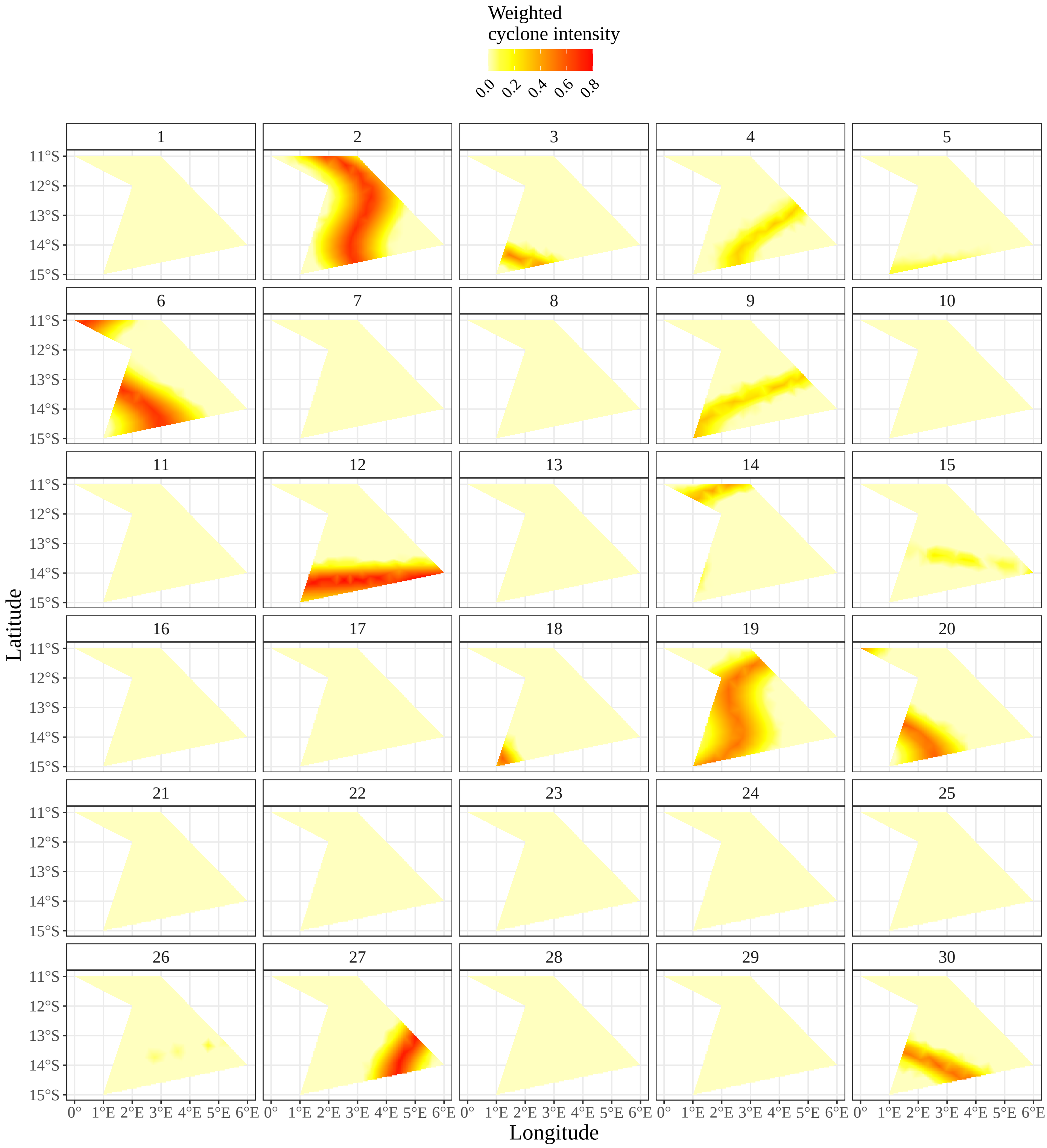

Figure 6: Weighted relative intensity of cyclone in space and time for the 30 years.

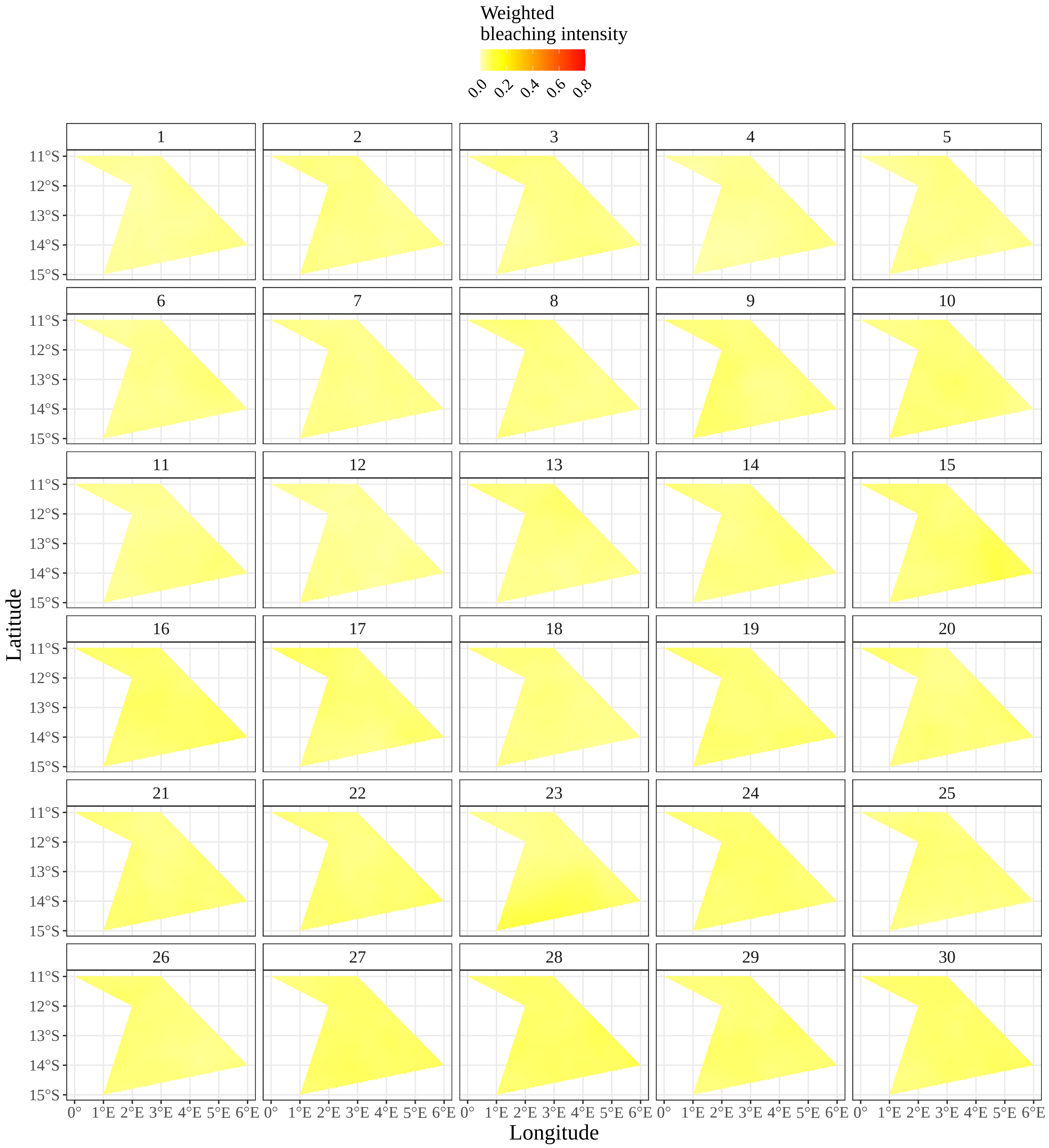

Figure 7: Weighted relative intensity of heat stress or bleaching in space and time for the 30 years.

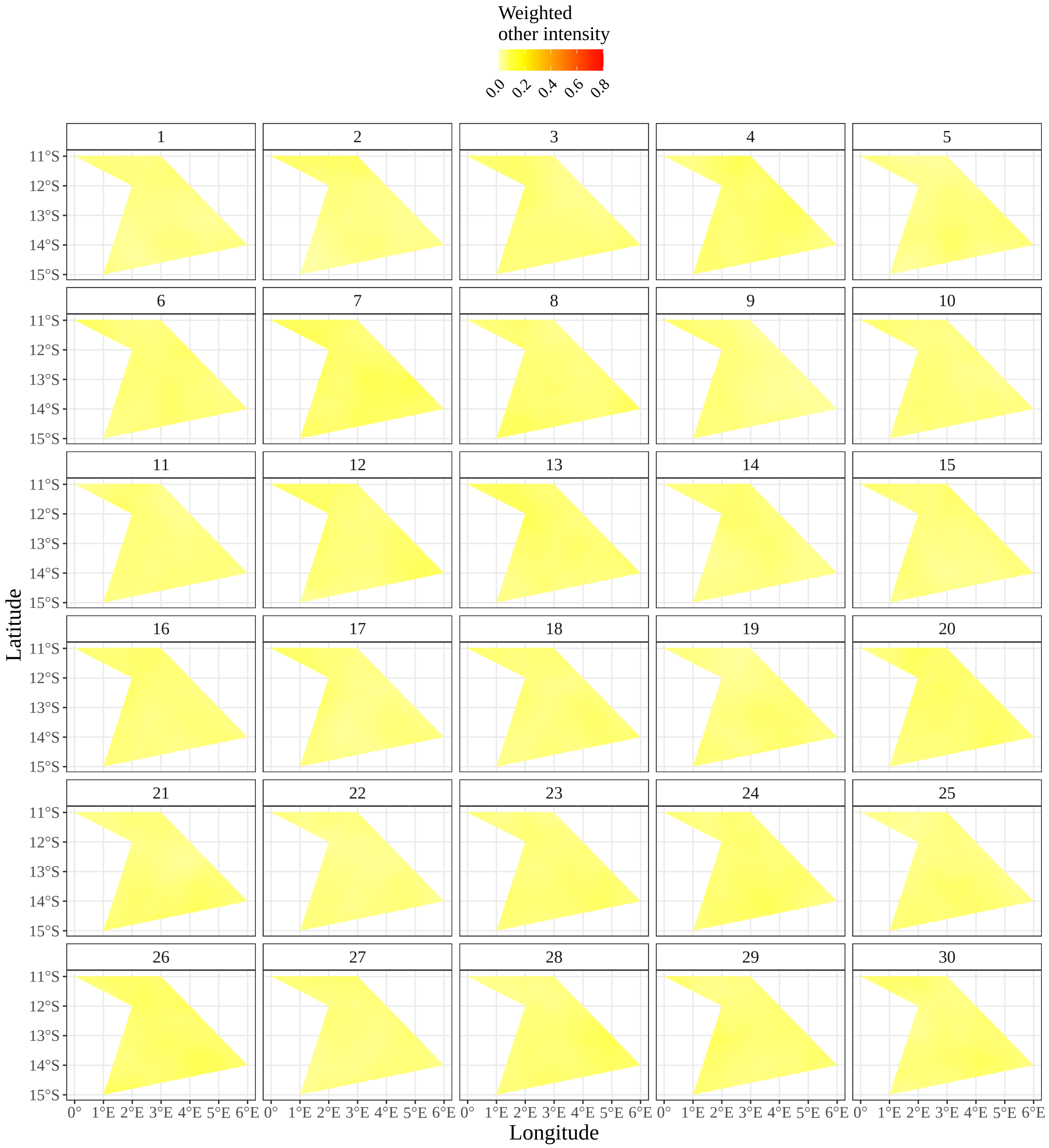

Figure 8: Weighted relative intensity of other disturbances in space and time for the 30 years.

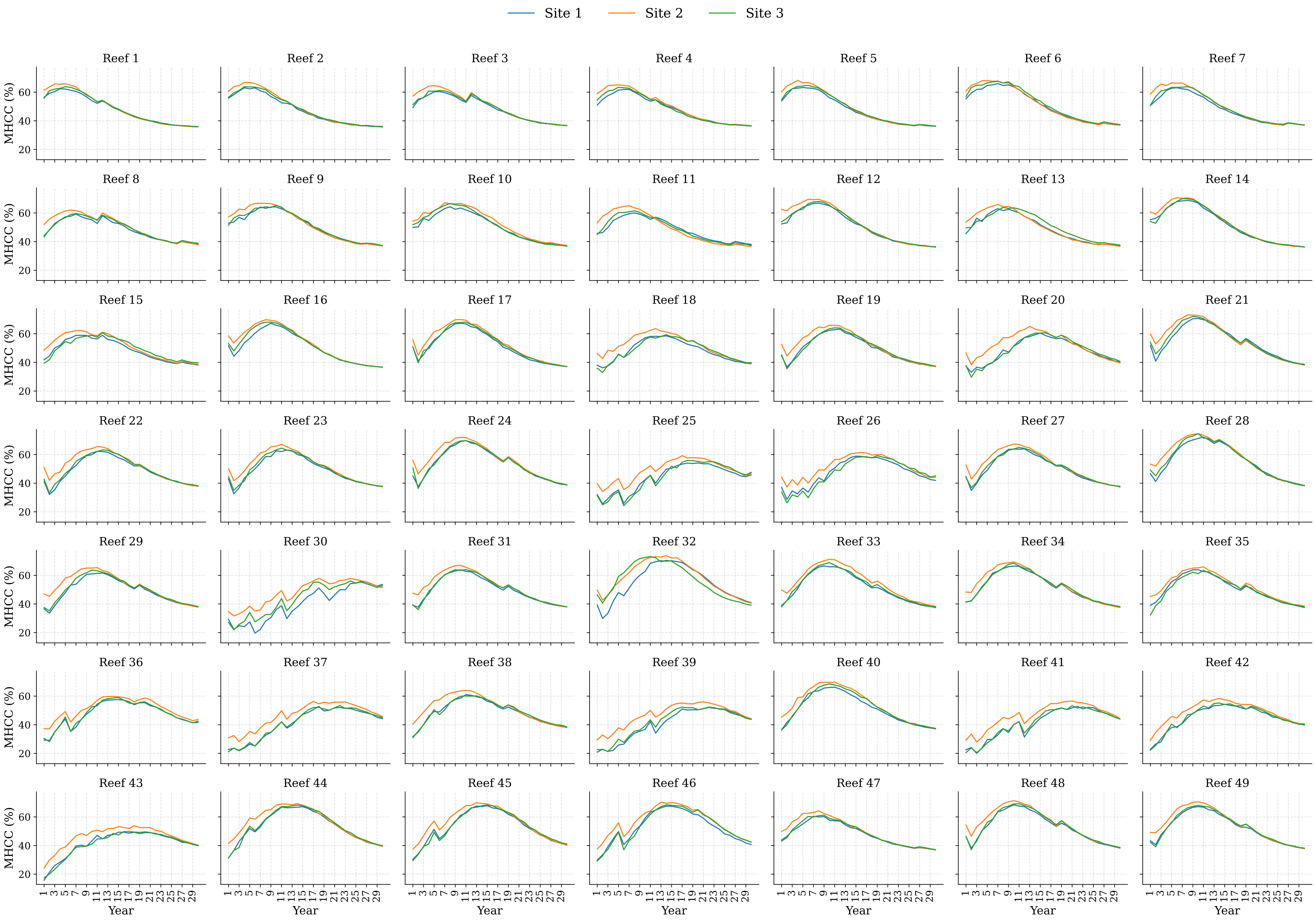

Figure 9: Time series of the observed MHCC at 10m depth. There are three sites for each reef and each line represents a site.

#### 2.2 Modelling frameworks

##### 2.2.1 Generalised additive model

Generalised linear models (GLMs) allow non-normal error distributions, additive terms, and non-linear relationships (Beery et al., 2021) with limited ability to deal with non-linear relationships (Hui et al., 2022). For a GLM, the response model is $Y \sim P(\mu)$, where $P$ is an exponential family distribution with $\mathbb{E}[Y] = \mu$ (Wood, 2017). The mean is modelled by, $g(\mu) = \alpha + \sum_{j=1}^{p} \beta_j x_j$ of $p > 1$ features and $g(.)$ is the link function that transforms the constrained $\mu$ to the real line. The link function can be chosen flexibly based on the distribution of the response variable (Molnar, 2019). GLM expresses the relationship between the mean response $g(\mu)$ and the predictor variables $x_j$ by Equation 4, where $\alpha$ is an intercept term and $\beta_j$ are the parametric terms.

$$g(\mu) = \alpha + \sum_{j=1}^{p} \beta_j x_j \tag{4}$$

While GLMs are non-linear in the sense that the link function is a non-linear transformation when the response is non-Gaussian, GAMs provide a further generalisation though replacing the linear terms $\beta_j x_j$ with functions $f_j(x_j)$ which are called smooth functions of the predictor variables (Simpson, 2018).

##### 2.2.2 Random forest

In a random forest (RF) predictions are calculated by taking the ensemble average of the $B$ independent regression trees. That is, we calculate the RF prediction from total number of trees ($B$) is given by $\hat{f}(x)$, where $f_b(x)$ is the prediction relevant to each individual tree with the input $x$ (a vector of features) (Lei et al., 2020), according to,

$$\hat{f}(x) = \frac{1}{B} \sum_{t=1}^{B} f_b(x). \tag{5}$$

**Algorithm 1** Random Forest

1. For $b = 1$ to $B$ do:
   - 1.1. Draw a random sample of size $N$ with replacement (bootstrap) from the training data.
   - 1.2. until the minimum node size is reached do
     - Randomly select a subset of $m$ predictor variables from total $p$.
     - Pick the best predictor optimizes splitting criterion among the $m$.
     - Split the node into two child nodes.
2. Output the ensemble of trees.

   $\hat{f}(x) = \frac{1}{B} \sum_{b=1}^{B} f_b(x)$.

##### 2.2.3 Boosted regression tree

Algorithm 2 represents the algorithm for gradient boosting algorithm (Friedman, 2001). While boosted regression tree (BRT) produces ensemble model by boosting the loss function, it is also able to deal with multiple forms of loss functions, such as Gaussian, Laplace, and quantile regression (QR). Given $Y$ is the dependent variable and $X$ is the predictor space, model starts with $F_0(x)$ which is the initial constant value, and then trees are added iteratively to minimise the loss function, $L$. At each stage, a decision tree is chosen in such a way that it minimizes the loss given the current model $F_{m-1}$ and its fit $F_{m-1}(x_i)$ (Shaziayani et al., 2021). $F_M(x)$ represents the output of the cumulative model or the final ensemble once $M$ trees are added iteratively.

**Algorithm 2** Gradient Boosting

**Input:**

- a training set $\{(x_i, y_i)\}_{i=1}^{n}$
- a differentiable loss function $L(y, \hat{y})$
- number of boosting iterations $M$
- a learning rate $\nu$

1. Initialize the model with a constant value:

$$F_0(x) = \arg\min_{\gamma} \sum_{i=1}^{n} L(y_i, \gamma)$$

2. **for** $m = 1$ **to** $M$ **do**

- Compute the pseudo-residuals:

$$\tilde{y}_{im} = -\frac{\partial L(y_i, F_{m-1}(x_i))}{\partial F_{m-1}(x_i)}, \quad i = 1, \ldots, n$$

- Fit a regression tree to the $\tilde{y}_{im}$ values
- Let the terminal regions of the tree be $R_j$ for $j = 1, \ldots, J_m$
- For $j = 1, \ldots, J_m$, compute:

$$\gamma_{jm} = \arg\min_{\gamma} \sum_{x \in R_j} L(y_i, F_{m-1}(x) + \gamma)$$

- Update the model:

$$F_m(x) = F_{m-1}(x) + \nu \sum_{j=1}^{J_m} \gamma_{jm} 1\{x \in R_j\}$$

3: Output the final ensemble $F_M(x)$

##### 2.2.4 Artificial neural network

Depending on the complexity of the artificial neural network (ANN), the number of hidden layers and the number of neurons in each hidden layer can be determined (Park and Lek, 2016). As shown in Figure 10, there are many input signals $[X = (x_1, x_2, \ldots, x_n)]$ to neurons while each input has a corresponding relative weight $[W = (w_1, w_2, \ldots, w_n)]$ which is an adaptive coefficient that determines the intensity of the input signal. As a flexible tool, with sufficient data and proper initialisation, ANNs can provide optimal solutions by adjusting the inner structure. Introducing non-linearity into the output of a neuron is done by the activation function $f$, and $b$ is called the bias associated with the neuron. Then, the output of a neuron $y$ can be expressed as the equation given in Figure 10.

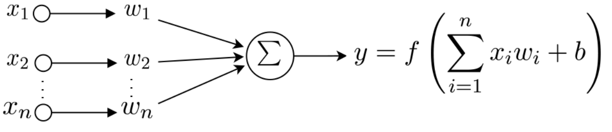

Figure 10: A neuron, where inputs are $x_1, x_2, \ldots, x_n$ and their corresponding weights are $w_1, w_2, \ldots, w_n$. $b$ is the bias associated with the neuron and activation function $f$ is applied to the weighted sum of the inputs.

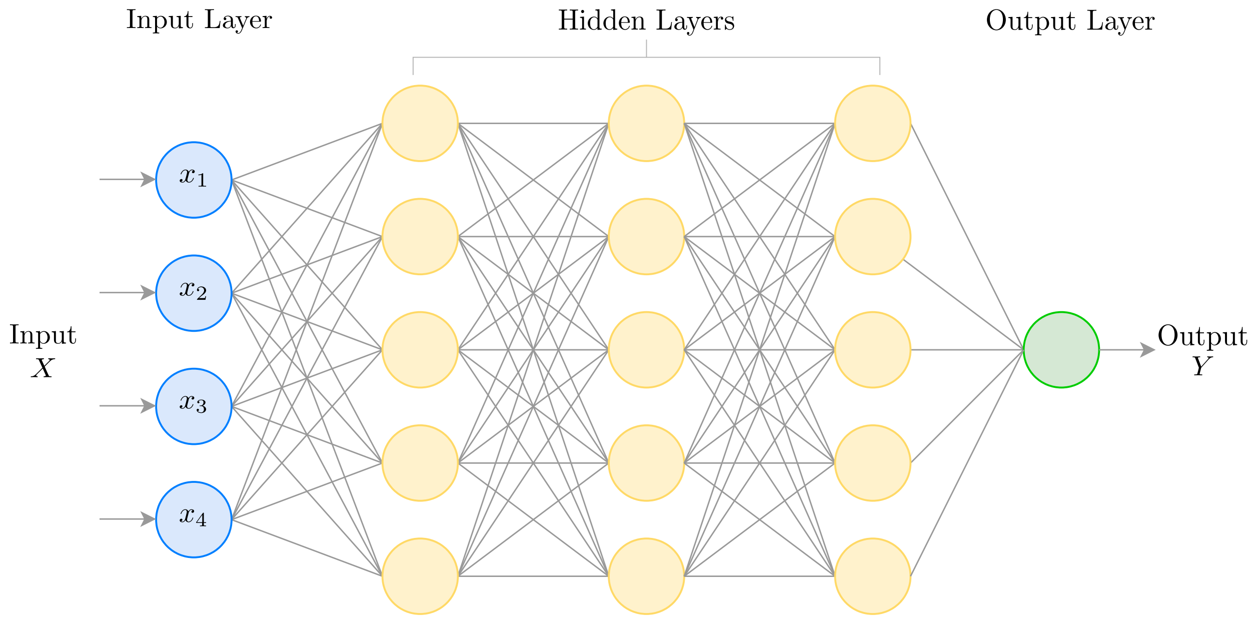

Figure 11: Multilayer Perceptron (MLP) with three hidden layers with five neurons in each hidden layer.

Multi layer perceptron (MLP) is defined as a fully connected ANN, as it consists of a system of interconnected neurons in each layer. The output of a neuron is adjusted by the connecting weight and fedforward to be an input to the subsequent layer's neurons. The architecture of a MLP can consist of several hidden layers of neurons. During training, the error or the difference between the desired and actual output is used to determine the appropriate adjustments should be made for the weights in the network to minimise the overall error of the MLP (Gardner and Dorling, 1998). This process is iteratively taken place and is known as Back-propagation (BP; Zhang and Li, 2017). MLP is also a feedforward ANN, as signals are transmitted in one direction, from input to output. Any non-linear function, except polynomial functions can be used as activation function and an appropriate combination of connecting weights and activation functions may lead to approximate any smooth, measurable function, allowing MLP to perform as an universal approximator (Popescu et al., 2009).

#### 2.3 Model construction and evaluation

In this work, all three ML models are tuned, involving a range of hyperparameters combinations. When tuning RF and BRT, we consider a range of tree complexity parameters such as number of estimators (trees), number of features to consider at every split, maximum tree depth, and minimum samples required to split a node. Additionally, for BRT we also consider tuning over a range of learning rates. MLP is tuned by exploring possible combinations of upto three hidden layers, with different number of neurons per each layer. The activation function, solver, regularization, number of iterations, and batch size are the other key parameters involved in the tuning process of MLP. When tuning ML models, random search is chosen over grid search since it is recommended as a computationally efficient method for hyperparameter optimisation, providing models with similar or better predictive performance compared to grid search (Bergstra and Bengio, 2012).

To evaluate the predictive performance of the models, we use a range of standard error metrics (Eq 6–9), where $y_i$ and $\hat{y}_i$ denote observed and predicted MHCC for a site in a given year, respectively. $n$ indicates the number of observations. $\bar{y}$ is the mean of the observed MHCC values.

$$MAPE = \frac{1}{n}\sum_{i=1}^{n}\left|\frac{y_i - \hat{y}_i}{y_i}\right| \times 100\% \tag{6}$$

$$RMSE = \sqrt{\frac{1}{n}\sum_{i=1}^{n}\left(y_i - \hat{y}_i\right)^2} \tag{7}$$

$$MAE = \frac{1}{n}\sum_{i=1}^{n}|y_i - \hat{y}_i| \tag{8}$$

$$R^2 = 1 - \frac{\sum_{i=1}^{n}(y_i - \hat{y}_i)^2}{\sum_{i=1}^{n}(y_i - \bar{y})^2} \tag{9}$$

### 3 Additional information for Section 4

#### 3.1 Model fit and predictive performance

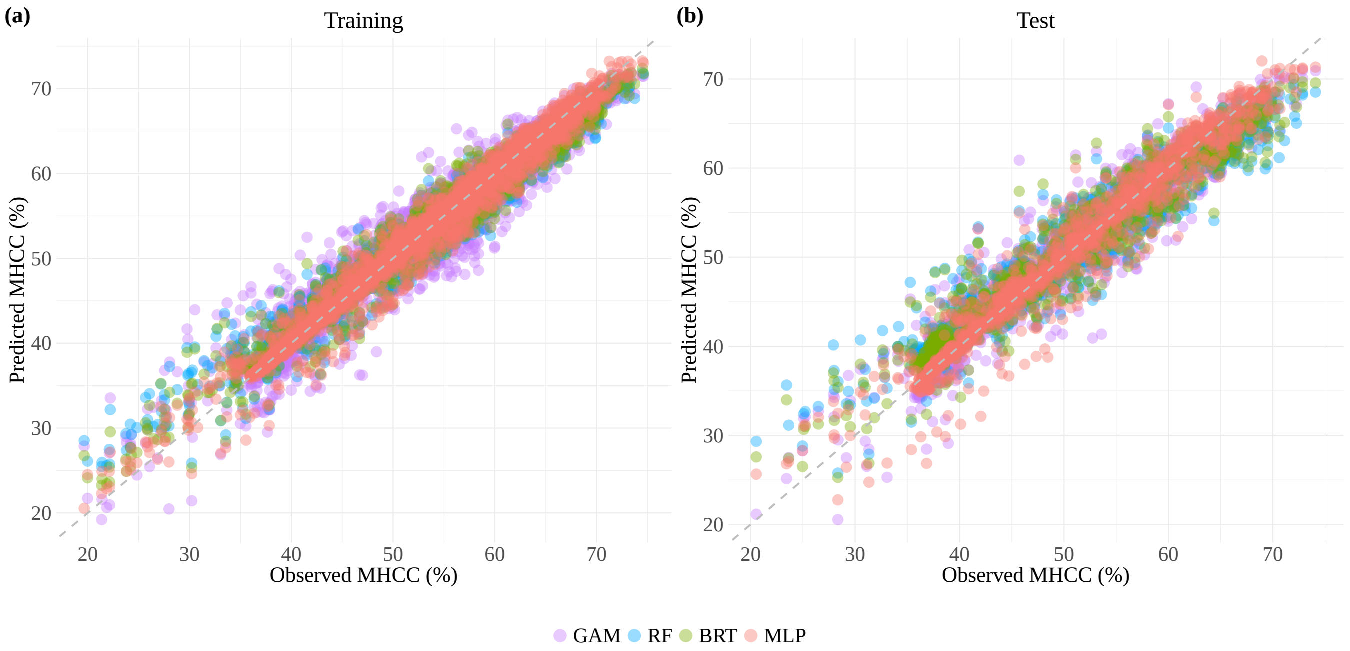

Figure 12: Predictive performance of the considered models for (a) withing-sample (training) data and (b) out-of-sample (test) data. y and x axes represent predicted MHCC by the models and corresponding observed value associated with the prediction, respectively.

As described in Section 3.2, we conduct a comparative analysis using three ML methods and a statistical model as a baseline model. Before evaluating explainability of these methods, their predictive performance is compared, and an overview of these models' performance is presented in Table 1.

Table 1: Summary of predictive model performance metrics, mean absolute percentage error (MAPE), root mean square error (RMSE), mean absolute error (MAE), and coefficient of determination ($R^2$) computed on predicted versus observed mean hard coral cover (MHCC) for within-sample (training) and out-of-sample (test) data. Bold values highlight the minimum values of MAPE, RMSE, MAE, and maximum of ($R^2$), under each error metric. Smaller values of MAPE, RMSE, and MAE indicates stronger predictive accuracy of the model. $R^2$ determines how well the variability of observed MHCC is explained by corresponding predicted values. While $R^2$ ranges from 0 to 1, higher values imply higher accuracy of predictions.

| | Within-sample (Training) | | | | Out-of-sample (Test) | | | |
|---|---|---|---|---|---|---|---|---|
| Model | MAPE | RMSE | MAE | $R^2$ | MAPE | RMSE | MAE | $R^2$ |
| GAM | 0.043 | 2.768 | 2.068 | 0.926 | 0.045 | 2.906 | 2.175 | 0.918 |
| RF | 0.027 | 1.870 | 1.307 | 0.966 | 0.047 | 2.923 | 2.187 | 0.917 |
| BRT | 0.024 | 1.719 | 1.206 | 0.971 | 0.041 | 2.654 | 1.959 | 0.931 |
| MLP | **0.021** | **1.540** | **1.050** | **0.977** | **0.030** | **2.243** | **1.463** | **0.951** |

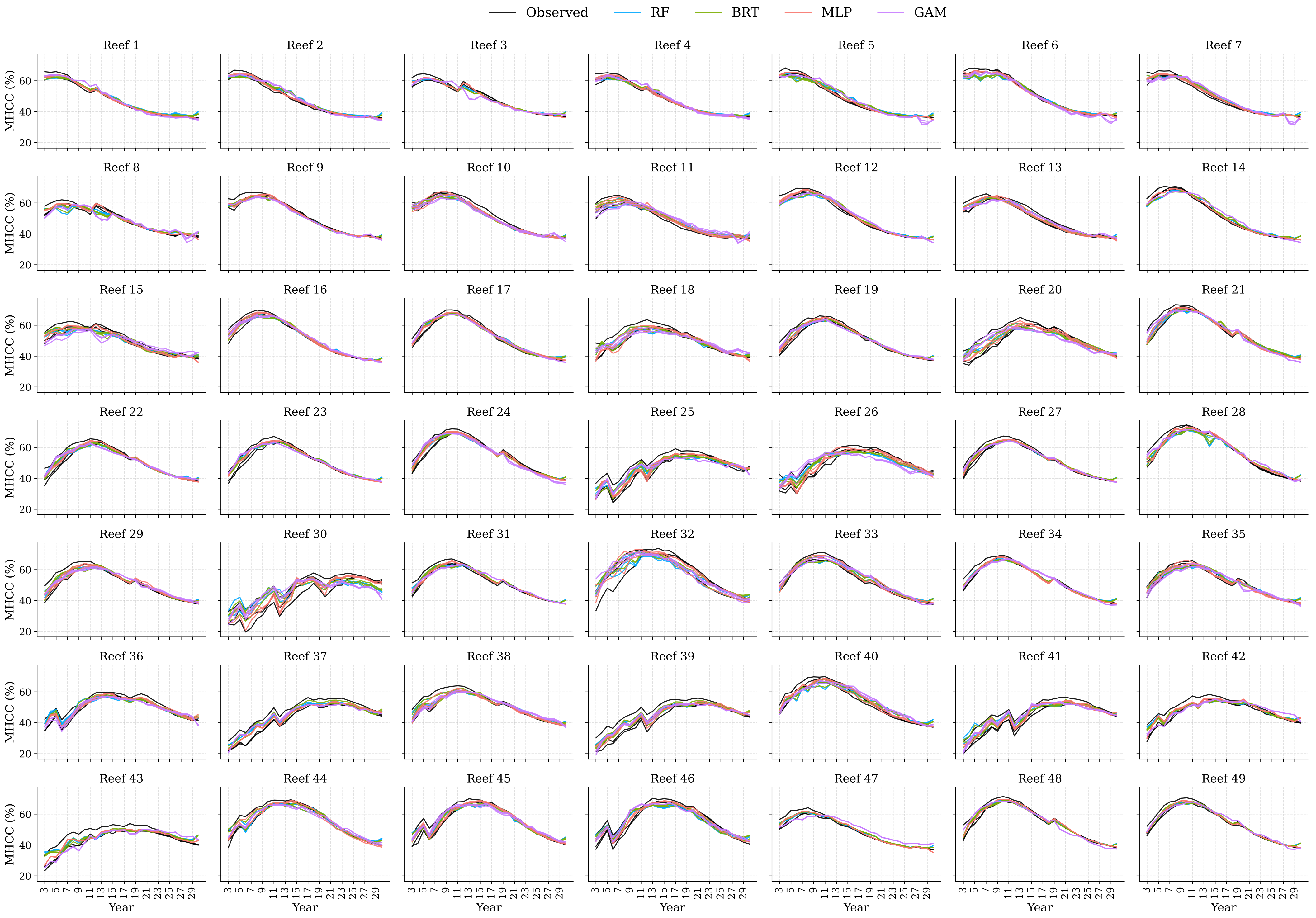
Observed
RF
BRT
MLP
GAM
Reef 1
Reef 2
Reef 3
Reef 4
Reef 5
Reef 6
Reef 7
Reef 8
Reef 9
Reef 10
Reef 11
Reef 12
Reef 13
Reef 14
Reef 15
Reef 16
Reef 17
Reef 18
Reef 19
Reef 20
Reef 21
Reef 22
Reef 23
Reef 24
Reef 25
Reef 26
Reef 27
Reef 28
Reef 29
Reef 30
Reef 31
Reef 32
Reef 33
Reef 34
Reef 35
Reef 36
Reef 37
Reef 38
Reef 39
Reef 40
Reef 41
Reef 42
Reef 43
Reef 44
Reef 45
Reef 46
Reef 47
Reef 48
Reef 49
MHCC (%)
20
40
60
Year
3 5 7 9 11 13 15 17 19 21 23 25 27 29

#### 3.2 Global explainability analysis

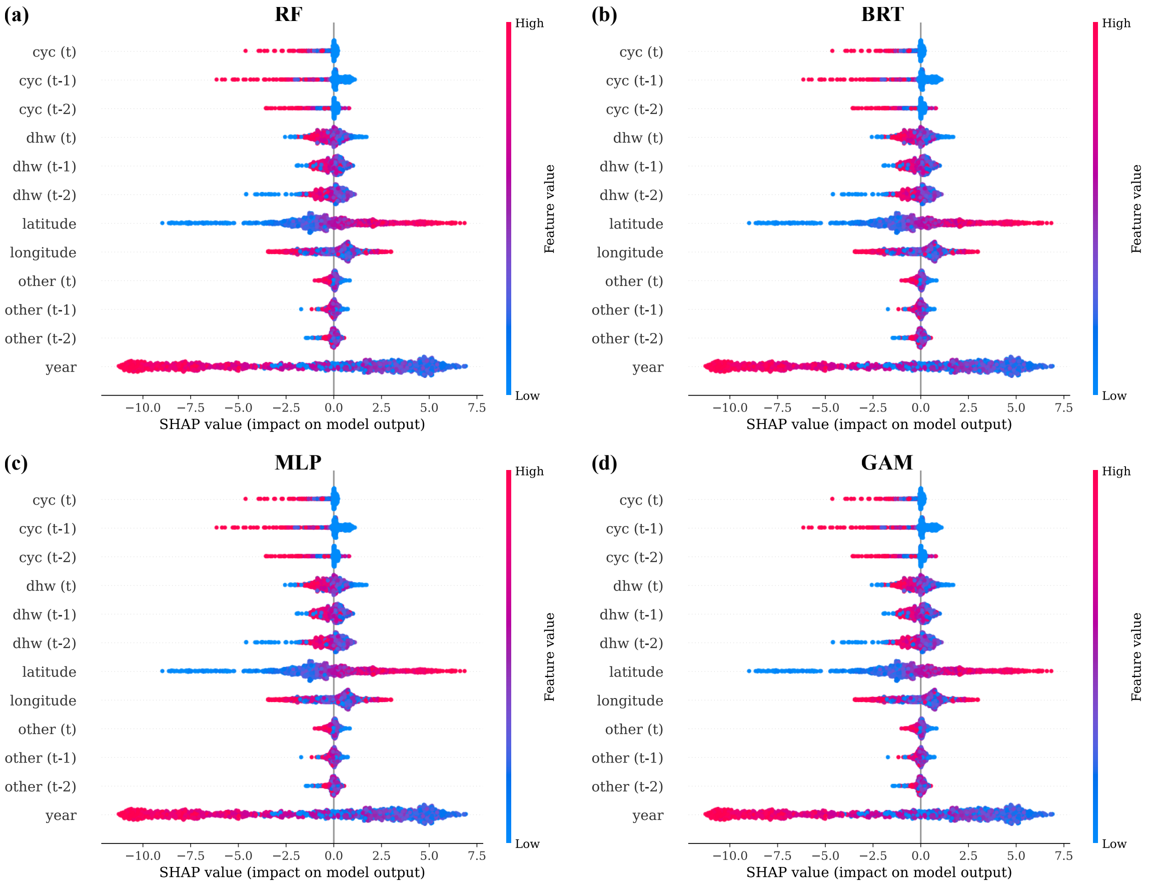

Figure 14: Beeswarm plot for (a) RF (b) BRT (c) MLP, and (d) GAM for the out-of-sample (test) data. The y-axis shows the features in the model and they are ordered alphabetically, for direct comparison, rather than the standard ordering by importance. The x-axis indicates SHAP value. Each dot represents a single site of a reef for a particular year. The absolute higher SHAP value indicates a higher effect on the model output or the prediction. The sign indicates the effect on the prediction, whether it is positive (increasing) or negative (decreasing). The magnitude of the SHAP values, expressed using a colour scale from red (high) to blue (low), is an indicator of how strong the effect of a feature is on the individual prediction of MHCC.

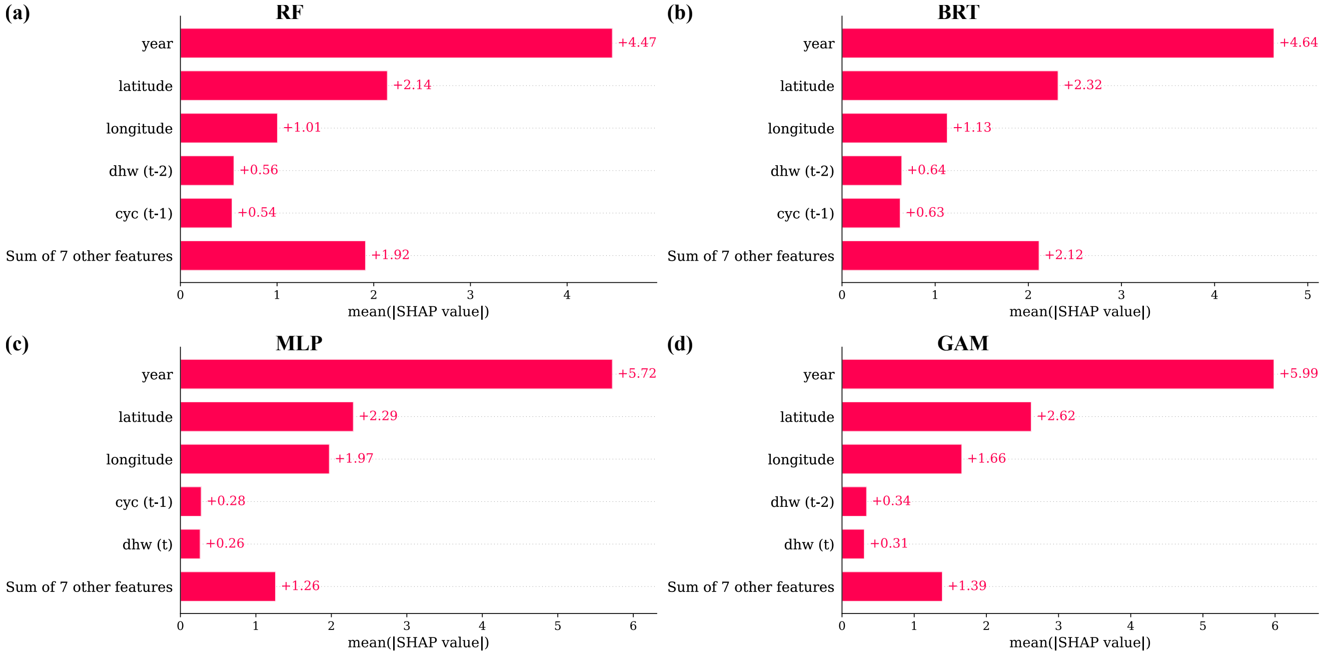

Figure 15: Mean SHAP for (a) RF (b) BRT (c) MLP, and (d) GAM for the within-sample (training) data. Mean SHAP values are obtained by averaging the absolute SHAP values for each feature across all the observations in the training data. Features with relatively larger mean SHAP indicate a higher effect on the predicted MHCC by the model.

#### 3.3 Representative scenarios of EDM

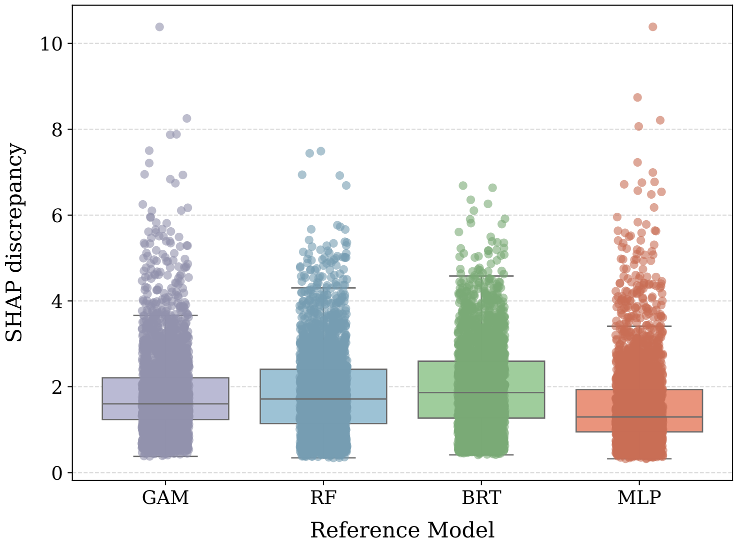

Figure 16: Boxplot of SHAP discrepancy for the training data. x-axis represents the reference model for the SHAP discrepancy measure.

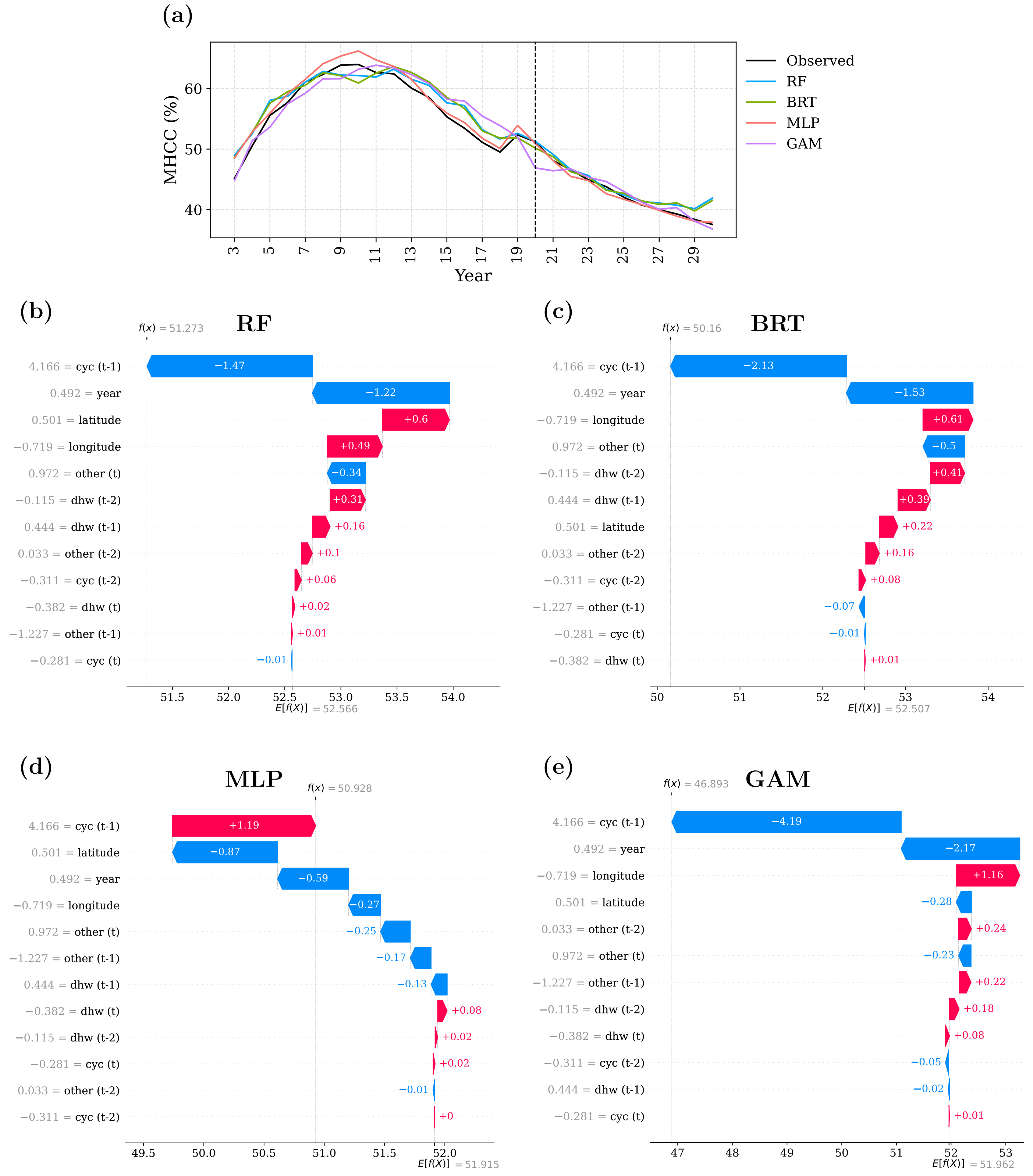

Figure 17: A high EDM common outlier ($E_1$–site 1 of reef 35, year 20). (a) shows the time series plot of observed and predicted values by the four models for site 1 of reef 35. The vertical dashed line represents the corresponding year of $E_1$. (b)-(e) are the waterfall plots from RF, BRT, MLP, and GAM models, respectively for this site in year 20. These waterfall plots visualise the individual contribution of features involved in the model to the final predicted MHCC, from the expected model output $E[f(x)]$ over the training dataset. The y-axis shows the features in the model, whereas the x-axis indicates SHAP value. The horizontal bars convey the direction of the contribution of the feature, whether it is positive (red) or negative (blue).

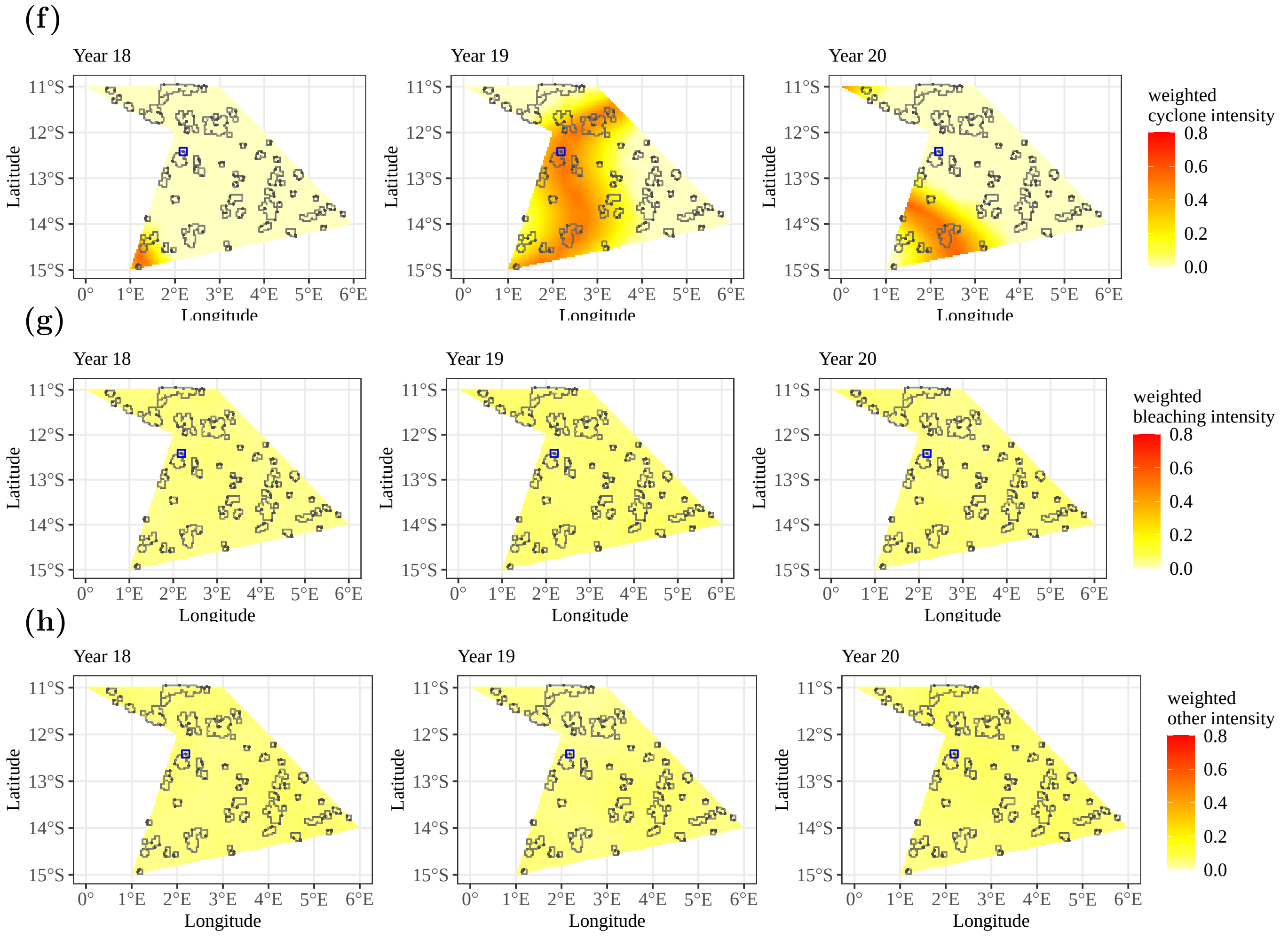

Figure 17: (continued) (f)–(h) show the relevant heatmaps (with 2 time lags) of weighted disturbances with the same colour scale. The grey coloured dots indicate the sites of each reef (irregular polygons) and, location of site 1 of reef 35 is highlighted from the blue coloured square.

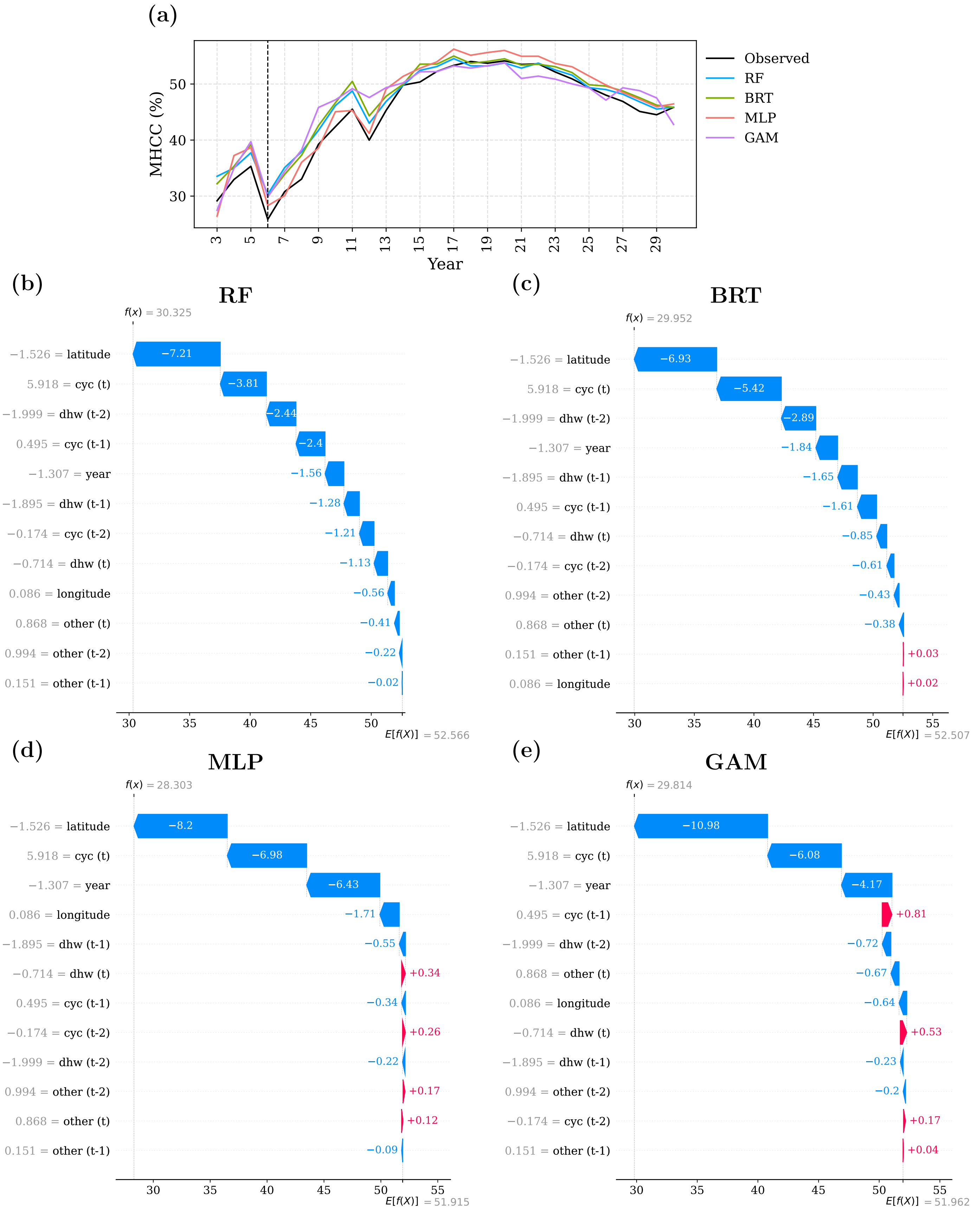

Figure 18: A clear cyclone event with low EDM ($E_2$–site 1 of reef 25, year 6). (a) shows the time series plot of observed and predicted values by the four models for site 1 of reef 25. The vertical dashed line represents corresponding year of $E_2$. (b)-(e) are the corresponding waterfall plots from RF, BRT, MLP, and GAM models, respectively for the this site in year 6. These waterfall plots visualise the individual contribution of features involved in the model to the final predicted MHCC, from the expected model output $E[f(x)]$ over the training dataset. The y-axis shows the features in the model, whereas the x-axis indicates SHAP value. The horizontal bars convey the direction of the contribution of the feature, whether it is positive (red) or negative (blue).

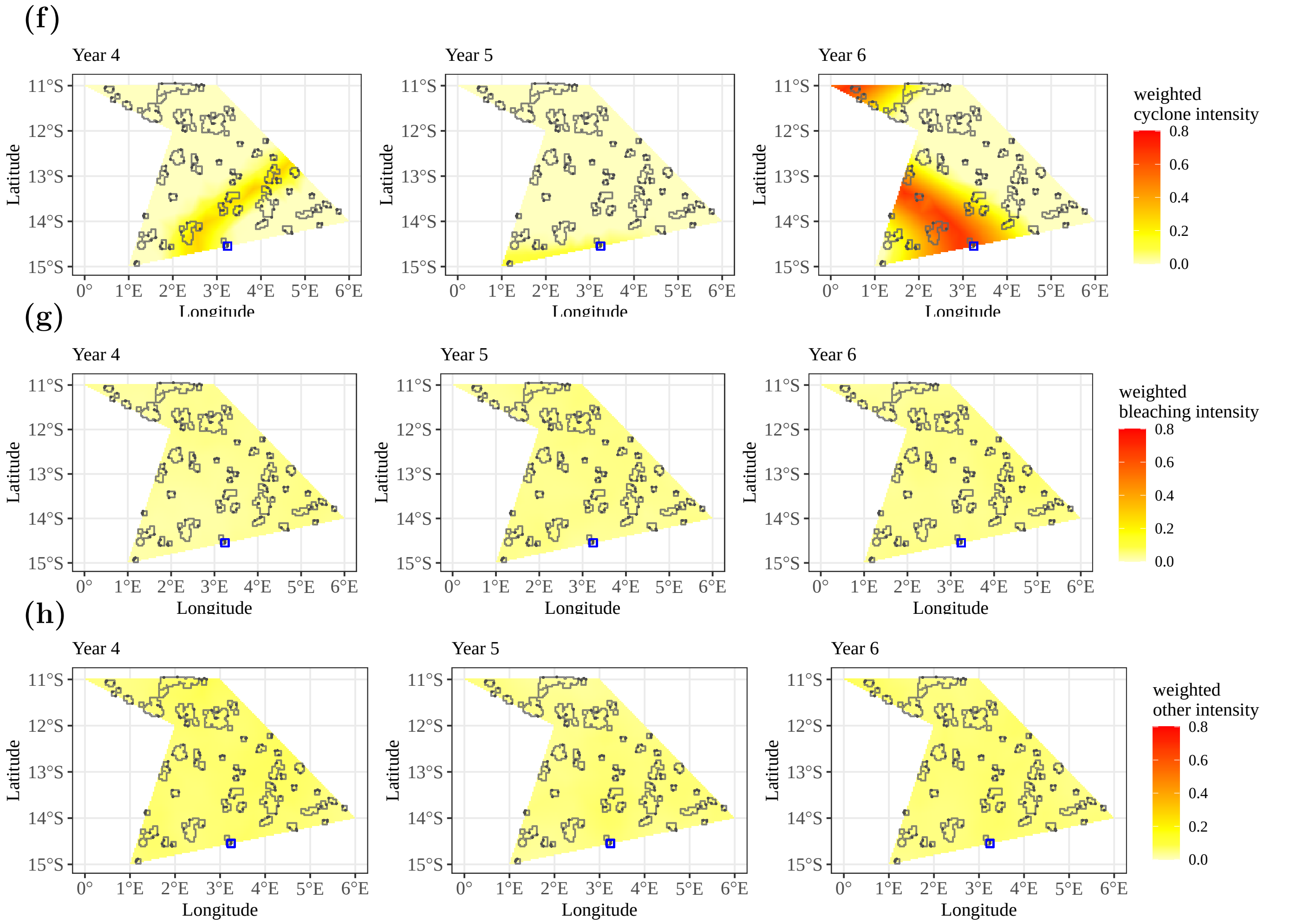

Figure 18: (continued) (f)–(h) show the relevant heatmaps (with 2 time lags) of weighted disturbances with the same colour scale. The grey coloured dots indicate the sites of each reef (irregular polygons) and, location of site 1 of reef 25 is highlighted from the blue coloured square.

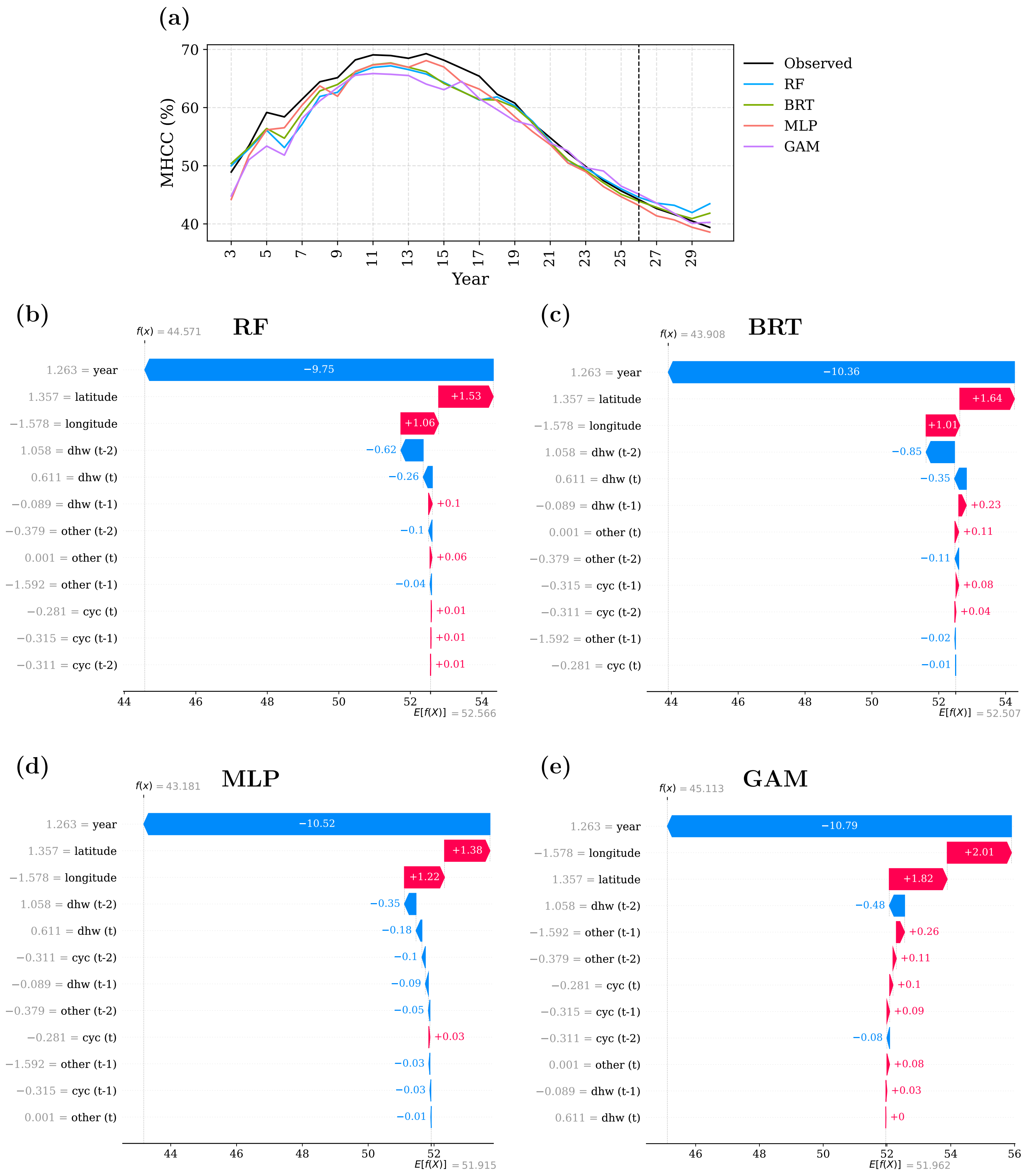

Figure 19: A low EDM point ($E_3$–site 2 of reef 44, year 26). (a) shows the time series plot of observed and predicted values by the four models for site 2 of reef 44. The vertical dashed line represents the corresponding year of $E_3$. (b)-(e) are the waterfall plots from RF, BRT, MLP, and GAM models, respectively for this site in year 26. These waterfall plots visualise the individual contribution of features involved in the model to the final predicted MHCC, from the expected model output $E[f(x)]$ over the training dataset. The y-axis shows the features in the model, whereas the x-axis indicates SHAP value. The horizontal bars convey the direction of the contribution of the feature, whether it is positive (red) or negative (blue).

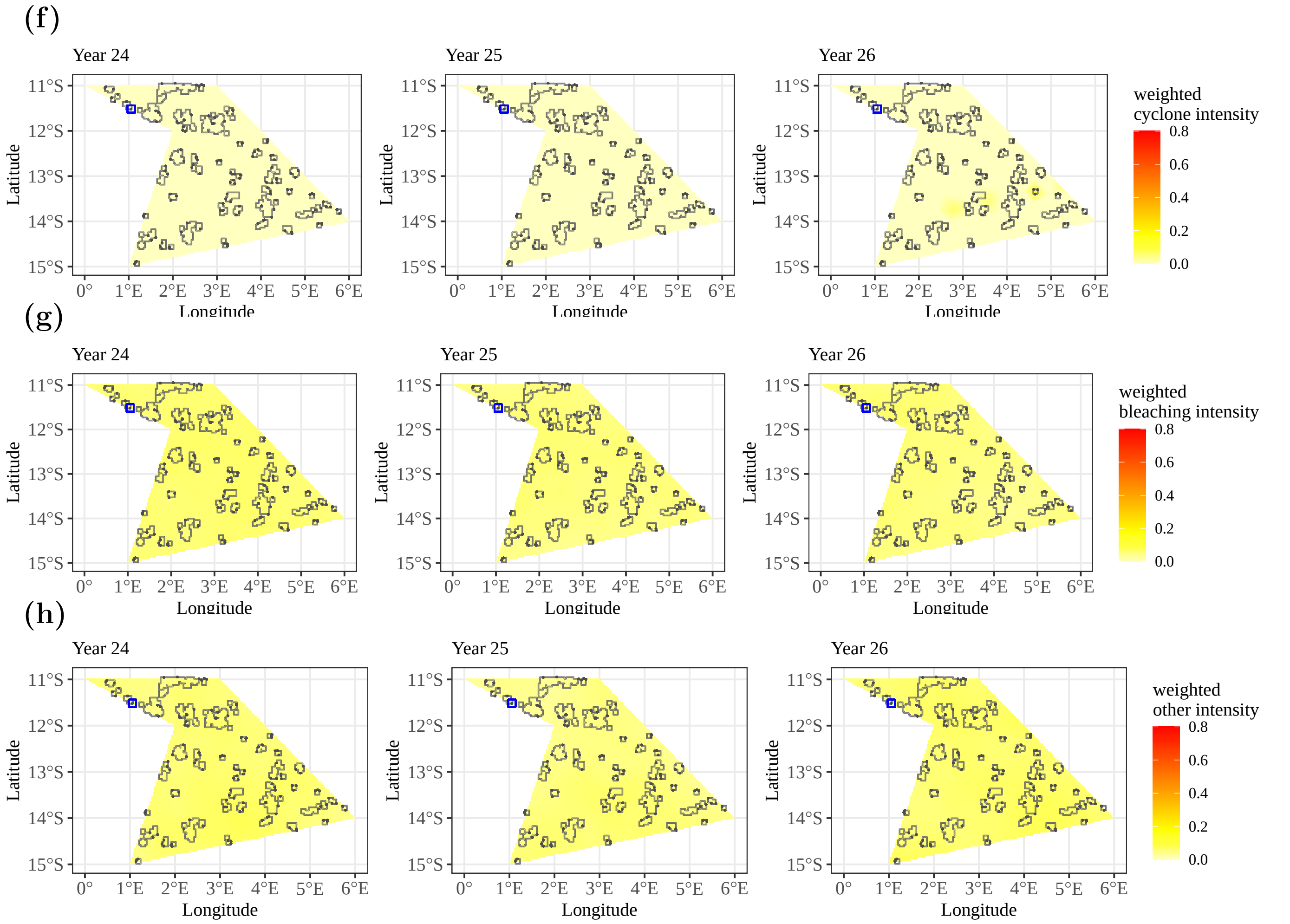

Figure 19: (continued) (f)–(h) show the relevant heatmaps (with 2 time lags) of weighted disturbances with the same colour scale. The grey coloured dots indicate the sites of each reef (irregular polygons) and, location of site 2 of reef 44 is highlighted from the blue coloured square.

##### 3.4 Prediction discrepancy

As given in Equation 10, $PD_{it}$, the prediction discrepancy for site $i$ at year $t$ is defined as the root mean squared difference between predictions from models and the observed value. In this equation, $\hat{y}_{itm}$ denotes the prediction from model $m$ for site $i$ at year $t$, while $y_{it}$ represents the observed value for site $i$ at year $t$, and $n$ indicates the total number of models.

$$PD_{it} = \sqrt{\frac{1}{n}\sum_{m=1}^{n}(\hat{y}_{itm} - y_{it})^2} \tag{10}$$

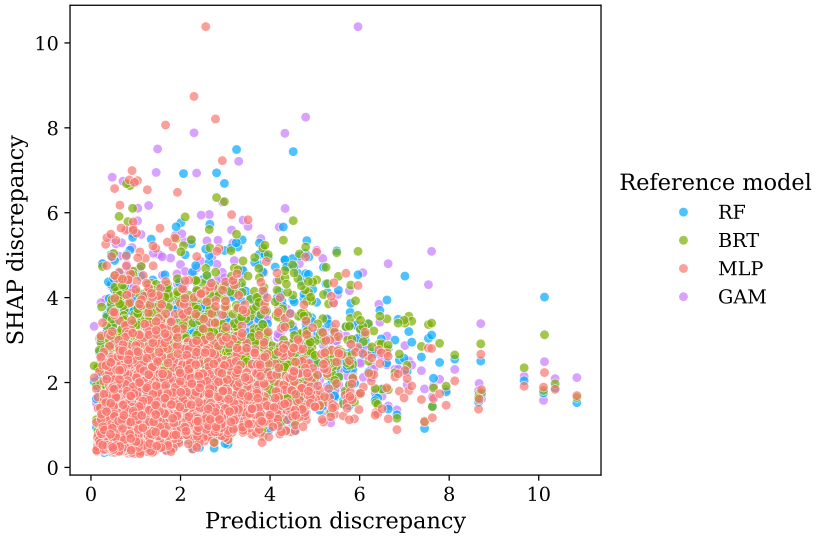

Figure 20: Scatterplot of SHAP explanation discrepancy versus prediction discrepancy. Prediction discrepancy is defined as the root mean squared difference between predictions from models and the observed value (Equation 10).
